# Spatial and functional mapping of the human pancreas reveals endocrine and exocrine cell states in health and metabolic disease

**DOI:** 10.64898/2026.06.30.735261

**Authors:** Yike Xie, Sandra Postic, David Pereyra, Roberta Ferrara, Alana Mullins, Johannes Pfabe, Mercedes Dalman, Kajsa Hallin, Sofie Ingvast, Nancy Smith, Mara Suleiman, Marta Tesi, Georg Gyoeri, Jule Dingfelder, Marta Gironella-Torrent, Patrick P. Starlinger, Lorella Marselli, Piero Marchetti, Patrick E. MacDonald, Marjan S. Rupnik, Olle Korsgren, Joan Camunas-Soler

## Abstract

The human pancreas contains diverse endocrine and exocrine cell populations whose spatial organization is essential for tissue physiology. While single-cell and spatial transcriptomics revealed molecular heterogeneity across pancreatic cell types, linking these states to physiological activity *in situ* has remained challenging. Here, we combined single-nucleus RNA sequencing, spatial transcriptomics, and functional calcium imaging across pancreatic samples in health and metabolic disease. We identified heterogeneous endocrine and exocrine cell states associated with obesity and diabetes, including inflammatory remodeling of acinar and ductal populations. To directly couple tissue physiology with molecular state, we developed Slice-seq, which integrates calcium imaging with spatial transcriptomics in acute pancreatic slices. Slice-seq linked local endocrine composition and transcriptional programs with β cell activity and identified extra-islet β cells with reduced glucose responsiveness and mitochondrial oxidative metabolism. Together, our study provides a framework for linking pancreatic cell states to tissue organization and physiological activity in health and disease.

## Introduction

From blood glucose regulation to digestive enzyme secretion, human pancreatic physiology relies on dynamic interactions among endocrine and exocrine cells and their tissue context. Single-cell and single-nucleus studies have revealed diverse cell subtypes and transcriptional states in the adult pancreas ^1,2^, some of them associated with diabetes and other pancreatic diseases. This diversity has been especially recognized among insulin-producing β cells ^3,4^, which also display variability in functional activity ^5^ and cytoarchitectural organization within islets ^6^. Studies in mouse and human islets have further established that local α, β, and δ cell interactions regulate hormone secretion and β cell activity through paracrine signaling and electrical coupling ^7–9^. In parallel, large-scale histological and whole-organ reconstruction analyses have revealed that human endocrine mass is broadly interspersed throughout the exocrine pancreas and frequently organized into small structures that do not conform to classical islet architecture ^10,11^. Despite the central role of β cell functional decline in diabetes ^12,13^, how this spatial heterogeneity relates to endocrine cell activity and dysfunction in disease remains unknown. Methods capable of integrating molecular states, tissue architecture, and physiological activity have the potential to address this gap. Whole-islet physiological studies have shown that islet composition is associated with secretory phenotypes and genetic risk for diabetes ^14^. At the single-cell level, Patch-Seq linked transcriptional heterogeneity with electrophysiological activity in isolated human α and β cells, identifying transcripts associated with dysfunction in diabetes ^15,16^. However, dissociation-based approaches inherently lack tissue context and local cellular interactions, making it difficult to map these findings onto islet and tissue cytoarchitecture. These limitations also impact understanding of exocrine physiology, as dissociation of enzyme-rich exocrine tissue remains technically challenging ^17,18^. Spatial transcriptomic approaches partially circumvent this limitation, enabling mapping of endocrine and exocrine heterogeneity without tissue dissociation ^19,20^. However, these methods currently lack direct information on dynamic patterns of cell activity.

Intracellular calcium signaling is a central regulator of stimulus-secretion coupling in electrically excitable endocrine cells through voltage-activated Ca^2+^ channels ^21,22^. In acinar and ductal cells, it also modulates fluid, electrolyte, and enzyme secretion through Ca^2+^ release from intracellular stores ^23,24^. Because Ca^2+^ dynamics can be monitored in tissue slices ^25,26^, high-resolution Ca^2+^ imaging in acute pancreatic slices provides a powerful approach to study cell activity in both mouse and human pancreas ^26–28^.

Here, we combined spatial transcriptomics, single-nucleus RNA sequencing, and functional Ca²⁺ imaging to map the human pancreas across molecular, spatial, and physiological dimensions. This multimodal approach enabled functional characterization of endocrine and exocrine cell types *in situ*, revealed a shift from mature secretory to inflammatory states in acinar and ductal cells during obesity and T2D, and linked islet cell composition to β cell Ca^2+^ activity. To directly interrogate the relationship between physiological activity and gene expression *in situ*, we developed Slice-seq, which combines sequential Ca^2+^ imaging and spatial transcriptomics in the same acute pancreatic slice. Slice-seq enabled direct investigation of gene expression and cell activity within the same tissue section, allowing characterization of small endocrine structures that do not conform to classical islet architecture and revealing altered metabolic coupling in extra-islet β cells. Overall, our study provides a comprehensive view of pancreatic organization across spatial scales and links physiological activity to gene expression within its native tissue context.

## Results

### A multimodal approach to map spatial, transcriptional, and functional features of the human pancreas

To investigate how endocrine and exocrine cell states vary across health and metabolic disease, we profiled human pancreatic tissue using two parallel workflows (**Fig. 1A)**. Fresh-frozen samples were used for single-nucleus RNA sequencing (snRNA-seq), immunohistochemistry, and spatial transcriptomics to define cell types, transcriptional programs, and tissue organization. In parallel, viable tissue slices were used for Ca^2+^ imaging followed by spatial transcriptomics (Slice-seq), enabling integration of functional activity with spatial gene expression while preserving local tissue architecture. In total, we profiled pancreatic tissue from 42 donors including non-diabetic lean (ND-Lean), non-diabetic obese (ND-Obese), pre-diabetic (Pre-T2D), type 2 diabetic (T2D), and type 1 diabetic (T1D) conditions (**Fig. 1B**; **Supp. Table 1**). Data were balanced across conditions, except in the ND-Obese group, which contained only females (**Supp. Table 1B)**. Across modalities, 34 samples were analysed by snRNA-seq and 28 by spatial transcriptomics across non-diabetic and diabetic conditions. In addition, pancreatic slices from 9 donors (primarily non-diabetic) underwent Ca^2+^ imaging, of which 5 were additionally profiled with Slice-seq.

**Figure 1.**
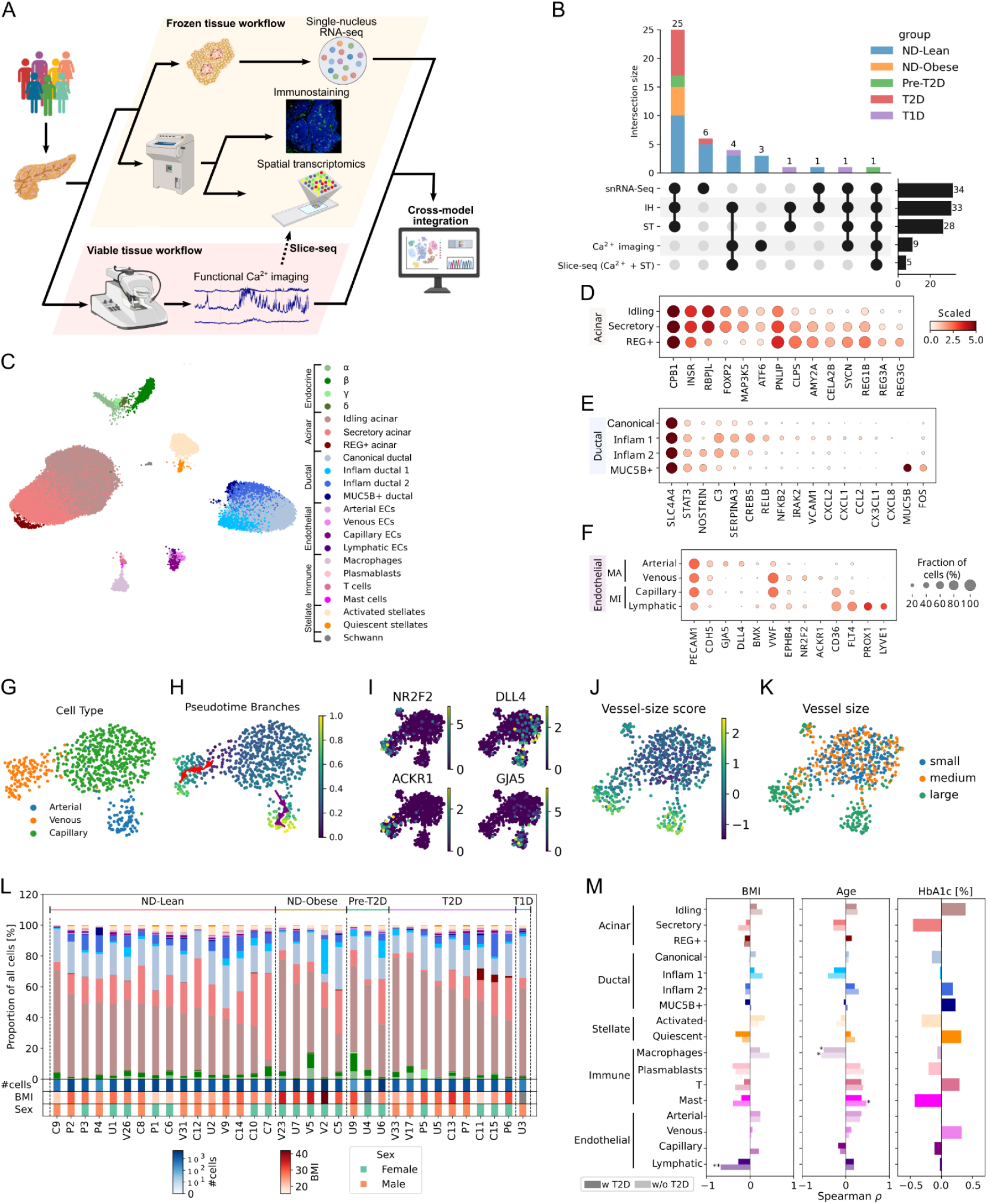
Experimental workflow and multimodal characterization of pancreatic cell states. **(A)** Study design. **(B)** UpSet plot showing donor overlap across profiling modalities. Stacked bar plots show the distribution of disease conditions across profiling technologies (top) and the total number of samples per method (right). **(C)** UMAP of 56,680 nuclei from snRNA-seq, colored by cell type and subtype. (**D**-**F**) Marker gene expression for acinar (D), ductal (E), and endothelial (F) cells. (**G-K**) Endothelial cell analysis: UMAP colored by cell type (G), pseudotime with inferred arterial (red) and venous (yellow) branches (H), zonation marker expression (I), vessel size score (J), and vessel size categories (K). **(L)** Relative abundance of cell subtype per sample based on color code from (C). **(M)** Correlation between subtype abundance (within cell type) and donor traits; lighter bars show analyses excluding T2D donors; darker bars show analyses including T2D donors (* p<0.05, ** p<0.01).

We first characterized cellular composition and transcriptional states using snRNA-seq, identifying all major pancreatic cell types with consistent representation across donors and clinical groups (**Fig. 1C**; **Supp. Fig. S1A-D)** and comparable quality metrics (**Supp. Fig. S1E-H)**. We identified multiple cell subtypes across acinar, ductal, and endothelial populations, and their transcriptional profiles were further characterized by pseudo-bulk differential expression analysis of snRNA-seq data (FDR<.1, **Supp. Table 2**).

Acinar cells comprised idling, secretory, and REG+ states, consistent with previous reports (**Fig. 1D**) ^29^. Ductal cells included canonical and MUC5B+ states (**Fig. 1E**) ^18^, along with two inflammatory subtypes enriched for complement system *C3* and *SERPINA3*, with one subtype exhibiting increased expression of NF-κB genes and multiple chemokines. These inflammatory subtypes may correspond to a recently described murine ductal population, enriched in medium-to-large ducts ^18^. Endothelial cells (ECs) spanned macrovascular (arterial and venous) and microvascular (capillary and lymphatic) populations (**Fig. 1F**). Diffusion-based pseudotime and branching analysis to ECs revealed two mirrored trajectories from capillary to arterial and venous states (**Fig. 1G,H**), with capillary cells expressing either arterial (*DLL4*, *GJA5*) or venous (*NR2F2*, *ACKR1*) markers (**Fig. 1I**). A vessel size score derived from pseudotime-associated genes identified a gradient from small to large vessels, with highest scores at the termini of macrovascular populations (**Fig. 1J,K**; **Supp. Fig. S1I**).

Relative abundances of cell types and subtypes in each sample were then quantified from the snRNA-seq data, with no major changes in composition across groups (**Fig. 1L**). We next examined associations between subtype abundance and donor traits, including age, BMI, and HbA1c. BMI was primarily associated with reduced abundance of lymphatic endothelial cells, whereas age was associated with immune cell subtype composition (**Fig. 1M**; **Supp. Fig. S1J-M**). In contrast, HbA1c did not show significant associations with cell type composition, consistent with previous islet studies showing preserved endocrine cell composition in T2D ^30^, and extending these findings to the exocrine pancreas.

### Shared inflammatory and metabolic transcriptional changes in exocrine cells during obesity and T2D

To investigate changes in tissue organization across metabolic conditions, we mapped cell types and subtypes identified by snRNA-seq onto spatial transcriptomics data (**Fig. 2A**). Spatial data quality was consistent across conditions (**Supp. Fig. S2A**) and snRNA-seq profiles from the same donors were used as a reference for cell type deconvolution to maximize concordance (**Methods**). Neighborhood enrichment analysis reflected expected tissue organization, including islet, ductal, and vascular structures, as well as spatial proximity between stromal (PSCs) and ECs consistent with known perivascular interactions (**Fig. 2B**) ^31,32^. Notably, we observed reduced spatial coherence of islets and ducts in ND-Obese donors (**Fig. 2C**), with similar but less pronounced effects in T2D, suggesting altered pancreatic tissue organization (**Supp. Fig. S2B-C**).

**Figure 2.**
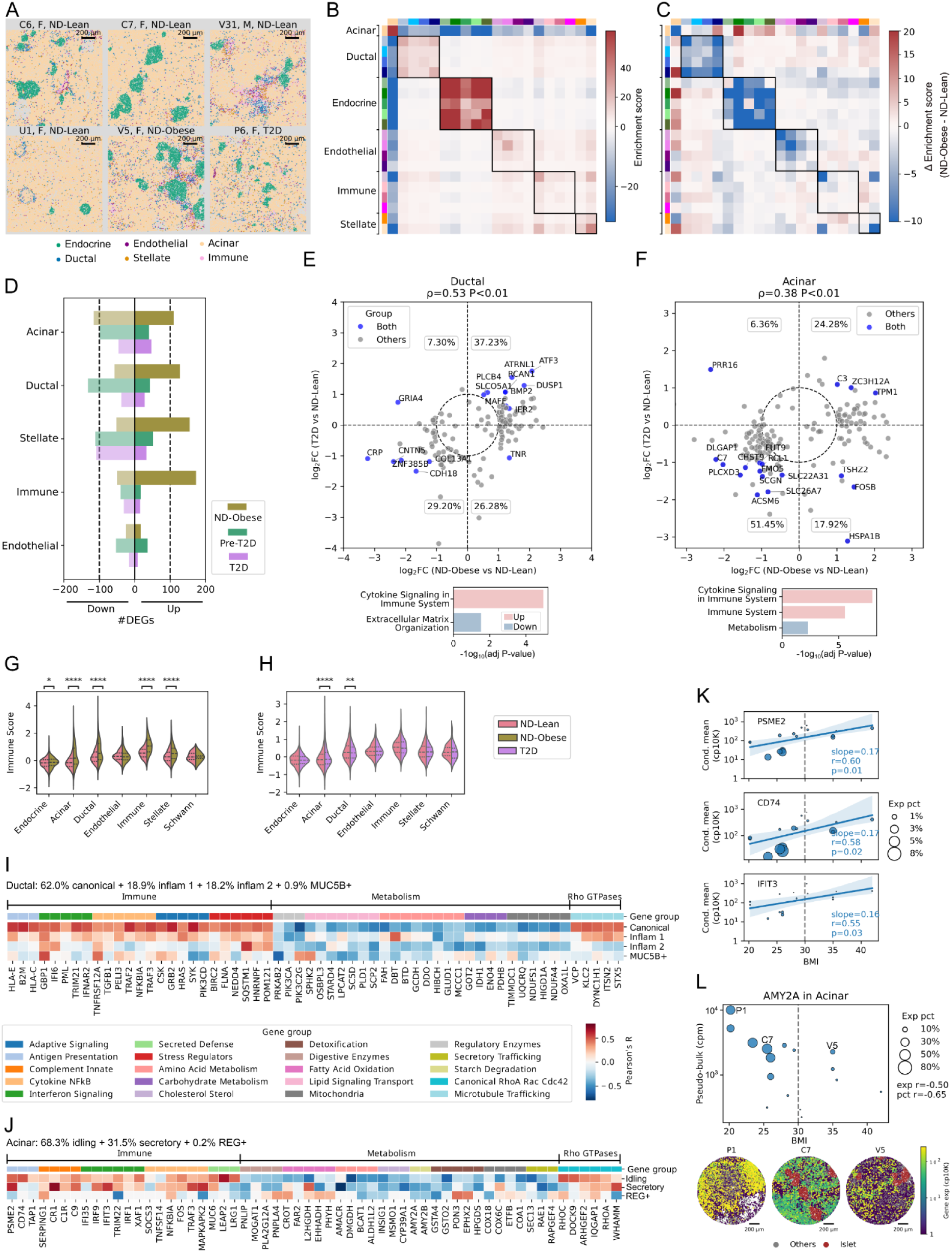
Spatial and transcriptional remodeling of exocrine cell types in obesity and T2D. (**A**) Representative spatial transcriptomic images from ND-Lean, ND-Obese, and T2D donors. Each spot is colored according to its cell-type assignment from Cell2location. (**B**) Neighborhood enrichment analysis showing expected spatial organization and colocalization of pancreatic cell subtypes in ND-Lean donors. (**C**) Difference in average neighbourhood enrichment score between ND-Obese and ND-Lean donors. (**D**) Number of differentially expressed genes in each cell type for ND-Obese, Pre-T2D, and T2D groups relative to ND-Lean. (**E**-**F**) Shared transcriptional changes in ductal (E) and acinar (F) cells in ND-Obese and T2D donors. Top: Log2 fold changes relative to ND-Lean controls for ND-Obese (x-axis) and T2D (y-axis) for genes identified in (D). Bottom: pathway enrichment analysis of concordantly upregulated and downregulated genes. (**G-H**) Enrichment scores of immune system genes across pancreatic cell types in ND-Obese (G) and T2D (H) donors relative to ND-Lean controls. (**I**-**J**) Pearson correlations between BMI and donor-level pseudobulk gene expression in ductal (I) and acinar (J) cell subtypes in non-diabetic donors. (**K**) Spatial transcriptomic expression of 3 representative BMI-associated immune genes in acinar cells, shown as average expression and fraction of expressing cells. (**L**) *AMY2A* expression in acinar cells as a function of BMI (top) and representative spatial expression patterns across donors (bottom). *Statistical significance was assessed using the Mann-Whitney test with Bonferroni-adjusted p-values (*p < 0.05, **p < 0.01, ***p < 0.001).

Next, we asked whether these alterations in tissue organization in obesity and T2D were associated with shifts in gene expression across cell types. Differential expression (DE) analysis revealed widespread changes across non-endocrine cell types, with the largest number of differentially expressed genes observed in ND-Obese donors compared to ND-Lean controls (**Fig. 2D**). Consistent results were obtained when restricting all obesity-related analyses to female donors **(Supp. Fig. S2E-J).** Differential expression was performed using memento, a single-cell specific method-of-moments approach, and validated using DESeq2 (**Supp. Fig. S2D)** ^33^. Acinar, ductal, stellate and immune cells showed the largest number of DE genes whereas ECs showed fewer differences across groups (**Fig. 2D**).

To determine whether obesity and T2D share common remodeling programs in exocrine cells, we next focused on acinar and ductal cells and compared transcriptional changes in both conditions. Analysis of DE genes in ductal cells (**Fig. 2E**), revealed shared patterns of gene expression between ND-Obese and T2D (rho=0.53, p<0.01), with most genes showing consistent directionality in fold-change (66.4%). A similar trend was observed for acinar cells (**Fig. 2F**), indicating partially shared transcriptional changes in obesity and T2D in these cell types. Shared changes in ductal cells included upregulation of stress and inflammatory regulators (*MAFF*, *ATF3*, *RCAN1*), cell signalling and remodelling factors (*BMP2*, *ATRNL1*), and downregulation of cell adhesion molecules (*CDH18*, *CRP*) (**Fig. 2E**). In acinar cells, ND-Obese and T2D donors showed upregulation of inflammatory and cytoskeletal-remodelling genes (*ZC3H12A*, *C3*, *TPM1*) and downregulation of transport, metabolic, and secretory genes (*SLC22A31*, *SCGN*, *ACSM6*, *FMO5*). Divergent changes included *FOSB*, *TSHZ2*, *HSPA1B* (up in ND-Obese, down in T2D) and *PRR16* (down in ND-Obese, up in T2D) (**Fig. 2F**).

Pathway analysis of genes showing concordant changes in ND-Obese and T2D donors (Quadrants 1 and 3 in **Fig. 2E-F**) showed upregulation of cytokine and immune signaling pathways in both ductal and acinar cells, with downregulation of extracellular matrix organization in ductal cells and metabolic programs in acinar cells (**Fig. 2E-F**, bottom panels). To determine whether these changes reflected global tissue or cell-type specific effects, we computed enrichment scores for immune signaling pathways across pancreatic compartments. In ND-Obese, we found significant enrichments in acinar, ductal, immune, and stellate cells (**Fig. 2G**), whereas in T2D only in acinar and ductal cells (**Fig. 2H**).

To further characterize transcriptional changes in acinar and ductal cells, we performed correlation analysis of gene expression with BMI across non-diabetic donors (**Fig. 2I-J)**. BMI was positively associated with immune and Rho GTPase signaling pathways, and negatively associated with metabolic pathways in both cell types. Exclusion of the highest-BMI donor (V2, BMI=42) led to reduced effect-sizes but did not significantly alter these results (**Supp. Fig. S2K-L**). Canonical ductal cells comprised 62% of the ductal pool and largely recapitulated global trends (**Fig. 2I**), showing positive correlations with MHC-I–based antigen presentation, interferon responses (*GBP1*, *IFI6*), NF-κB signaling (*NFKBIA*), stress response (*SQSTM1*), and negative associations with multiple metabolic pathways and mitochondrial respiration (*PDHB*, *ENO4*, *NDUFS1*). These associations were largely observed across ductal subtypes, except for Rho GTPase-related genes in MUC5B+ and Inflam 2 ductal cells. In acinar cells, most associations to BMI were driven by idling and secretory subtypes (**Fig. 2J**), including positive correlations with MHC-II antigen presentation, complement and innate defense, interferon signaling, and NF-κB signaling. We validated some of the top correlated genes (*PSME2*, *CD74*, *IFIT3*) in spatial transcriptomics, finding significant associations in this data modality as well (**Fig. 2K**). Conversely, BMI showed negative correlations with key pancreatic digestive enzymes (*AMY2A*, *AMY2B*, *PNLIP*), mitochondrial respiration, and secretory trafficking among other pathways. This trend was also confirmed by spatial transcriptomics data, with AMY2A showing the strongest association with BMI among digestive enzymes (**Fig. 2L**; **Supp. Fig. S2M**).

Altogether, these results indicate that obesity is associated with exocrine remodeling, characterized by a shift from metabolic programs towards inflammatory and stress-related transcriptional responses. In acinar and ductal cells, these changes partially overlap with those observed in T2D (**Fig. 2E-F)** and include reductions in mature secretory programs resembling those observed in early acinar remodeling ^34^.

### Macrophage infiltration and associations with islet cell gene expression in T2D

The upregulation of multiple IFN-γ-response genes in acinar (*IRF1*, *TRIM22*) and ductal (*GBP1*) cells in ND-Obese prompted us to investigate tissue-resident macrophages (see **Methods**). Surprisingly, macrophage abundance and activation state were not significantly associated with BMI, and enrichment of pro-inflammatory macrophages was only observed in the highest BMI donor (**Fig. 3A**; **Supp. Fig. S3A)** ^35,36^. This is consistent with recent multiplexed imaging studies in T2D reporting no increase in total immune cells (CD45+), macrophage abundance (IBA+) or activation in the exocrine pancreas ^37^. These observations suggest that the inflammatory and stress-response programs identified above in acinar and ductal cells may arise in the absence of local immune cell expansion.

**Figure 3.**
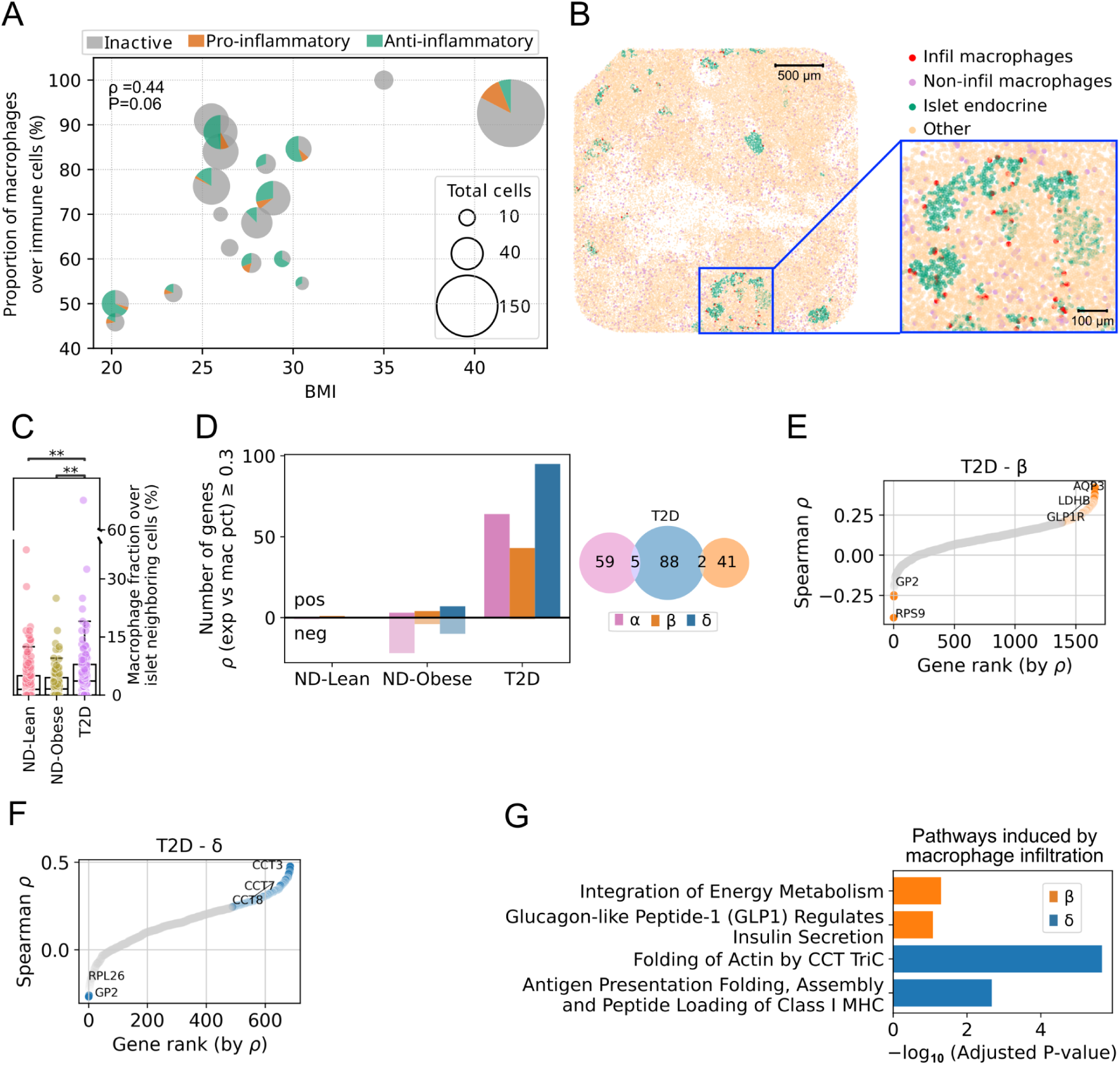
Macrophage infiltration and gene-expression changes in T2D islets. (**A**) Macrophage abundance and activation state in pancreatic tissue from ND donors as a function of BMI (snRNAseq data). Each pie chart represents a donor, Spearman’s correlation coefficient and p-value are indicated. (**B**) Representative spatial transcriptomic image showing infiltrating macrophages (V23). (**C**) Islet-associated macrophage abundance across donor groups, defined as the fraction of macrophages among all neighboring cells within 20 μm of islets. ** indicates p < 0.01 (Mann-Whitney test). (**D**) Number of genes correlated with islet-associated macrophage abundance in α, β, and δ cells (|ρ| > 0.3); Venn diagram shows gene overlap in T2D. (**E–F**) Ranked genes correlated with islet-associated macrophage abundance in β (E) and δ (F) cells from T2D donors. Representative genes highlighted. (**G**) Pathway enrichment analysis of genes positively associated with islet-associated macrophage abundance in β and δ cells in T2D donors.

In spatial transcriptomics, macrophages were sparsely detected and occasionally observed near islets (**Fig. 3B**; **Supp. Fig. S3B**). Islets from donors with T2D showed a significant increase in infiltrating macrophages, defined as those within 20 μm of an islet, which exhibited a higher pro-inflammatory signature than non-infiltrating ones (**Fig. 3C**; **Supp. Fig. S3C**). Within T2D islets, macrophage infiltration was associated with transcriptional changes across endocrine cell types (**Fig. 3D**), with the largest associations in δ cells (ρ>0.5 vs ∼0.3 in β cells) (**Fig. 3E-F**). Pathway analysis revealed increased expression of cytoskeletal remodeling genes in these δ cells (**Fig. 3G**; **Supp. Fig. S3D-E)**, suggesting alterations in δ cell state and potentially morphology.

Together, these findings indicate that macrophage infiltration and activation are unlikely to underlie the transcriptional changes observed in exocrine cells in obesity, which may instead reflect systemic or metabolic factors. In contrast, macrophage infiltration in T2D islets is associated with changes in endocrine cell states, especially in δ cells. Prior work has shown that δ cell morphology and cytoskeletal organization are closely linked to paracrine regulation within islets ^38–40^, raising the possibility that changes in δ cell state contribute to altered endocrine signaling in T2D.

### Islet heterogeneity and transcriptional changes in β-cell-rich islets

Recent 3D and large-scale histological analyses have shown that human islets are highly heterogeneous with many small endocrine clusters enriched in β-cells not conforming to classical islet architecture ^10,41^. However, the functional consequences of this heterogeneity are debated ^11,42^. Building on this, we used immunostaining to quantify islet morphology and endocrine cell composition, and spatial transcriptomics to ask whether β cell gene expression differs across islets of different composition.

We segmented 1670 individual islets from 23 donors, confirming previously reported heterogeneity in islet size, shape, and endocrine cell proportions (**Fig. 4A**; **Supp. Fig. S4A)** ^6,43^. Across donors, β cells were the most abundant endocrine population, followed by α and δ cells, with broadly comparable distributions across groups except for the expected reduction in β cell numbers in T1D (**Supp. Fig. S4B**). At the individual-islet level, β cell proportions also varied substantially, even within ND-Lean donors (IQR: 33.3 - 80.7**%**) (**Fig. 4B**), highlighting marked intra-donor islet heterogeneity.

**Figure 4.**
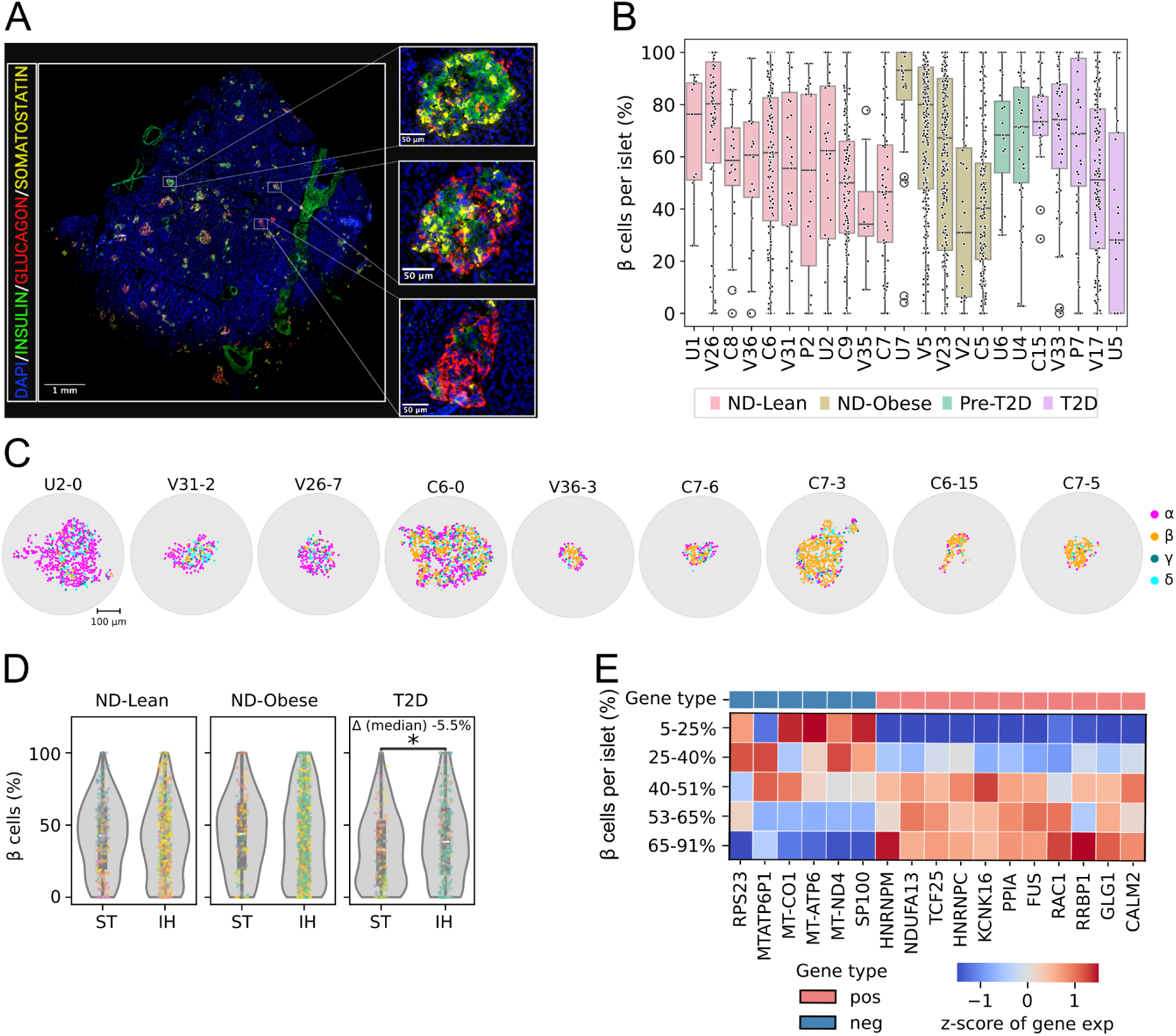
Heterogeneous β-cell composition and gene expression in human islets. (**A**) Immunostaining images of comparably sized islets with differing β cell abundance and endocrine cell composition. (**B**) Fraction of β cells per islet across donors based on immunostaining data. Each dot represents an islet. No significant differences were detected between groups by the Mann–Whitney U test. (**C**) Representative α-rich (left), mixed (middle), β-rich (right) islets identified by spatial transcriptomics. (**D**) Comparison of β-cell fractions per islet derived from spatial transcriptomics (ST) and immunostaining (IH) across donor groups. Each dot represents an islet. (**E**) Heatmap showing genes positively or negatively associated with the fraction of β-cells within individual islets in ND-Lean donors.

Equivalent heterogeneity in islet size, shape and composition was observed in spatial transcriptomics data (**Fig. 4C**), where islets were independently segmented using endocrine gene enrichment scores and spatial autocorrelation. Per-donor endocrine cell abundances were consistent across modalities (**Supp. Fig. S4C**), and spatial transcriptomics yielded per-islet β cell proportions consistent with immunostaining (**Fig. 4D**). To identify cell-type specific transcriptional programs associated with islet composition, we performed cell-type pseudobulk averages per islet, and correlated gene expression with % β cells in each islet. Simulated pseudobulk averages from snRNA-seq-data were used to derive a reference distribution to control for potential admixture effects (**Supp. Fig. S4D-E**). In ND-Lean donors, β-cell-rich islets showed higher expression of regulators of membrane excitability and calcium signaling (*KCNK16, CALM2*), and reduced expression of mitochondrial respiration genes (**Fig. 4E**). In humans, *KCNK16* encodes the islet-specific K⁺ channel TALK-1, which regulates β cell resting membrane potential and limits Ca^2+^ oscillations following glucose stimulation ^44,45^. Together, these results provide evidence that differences in islet composition are associated with transcriptional programs that support coordinated β cell activity and may underlie functional heterogeneity in human islets.

### Slice-seq links Ca^2+^ activity to cell types and gene expression programs in live tissue slices

To further investigate functional and transcriptional activity of human pancreatic cell types *in situ*, we developed Slice-seq, which integrates Ca^2+^ imaging and spatial transcriptomics in the same tissue slice (**Fig. 5A**). We established Slice-seq as a multi-step process: first, acute pancreatic slices (∼140 μm) are prepared and subjected to live Ca^2+^ imaging under controlled perfusion conditions ^25,26,28,46,47^; next, the same slice is carefully re-embedded in a flat OCT surface while preserving its orientation; finally, consecutive cryosections (∼10 μm) are collected by sectioning toward the imaged surface for spatial transcriptomics and immunostaining (see **Methods**). This approach generates flat, parallel sections aligned with the original slice axis, and enables the mapping of Ca^2+^ activity in the same islets that are subsequently profiled by immunohistochemistry for cell type annotation and with spatial transcriptomics for gene expression (**Fig. 5B**; **Supp. Fig. S5A**). Spatial transcriptomics was performed using Curio Seeker, which uses beads approximately the size of an islet cell to capture transcripts (∼10 μm). Although direct cell-to-cell mapping between modalities is limited by resolution and tissue deformation, Slice-seq enables spatial mapping of gene expression in close proximity to the imaged plane and within the same islets and pancreatic structures across modalities.

**Figure 5.**
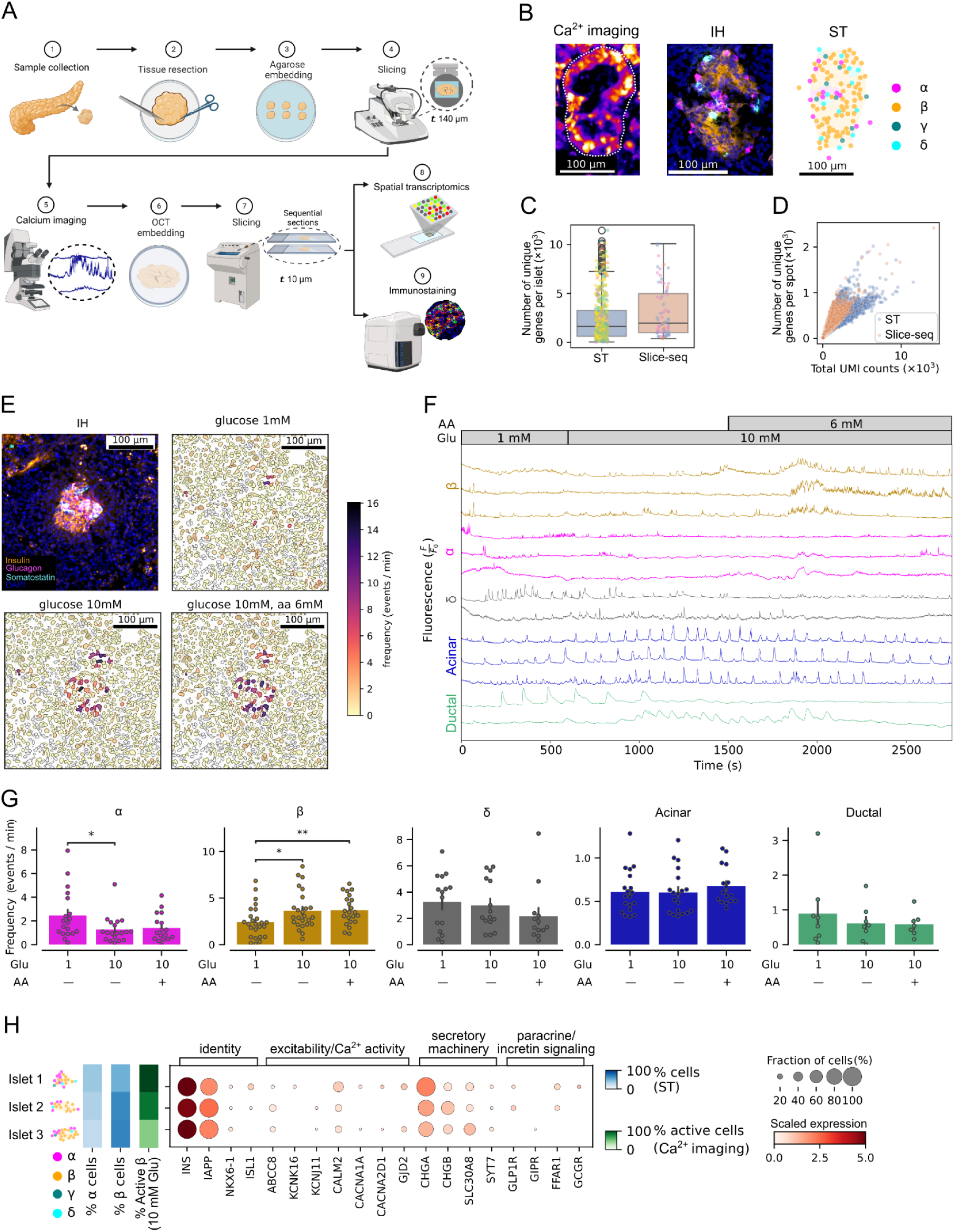
Slice-seq measurements in acute human pancreatic slices. (**A**) Workflow illustrating Slice-seq. (**B**) Ca^2+^ imaging (left; dashed lines indicate islet boundaries), immunostaining (middle), and spatial transcriptomics (right) form the same islet in a Slice-seq experiment. (**C**) Number of genes per islet in spatial transcriptomics and Slice-seq data. Each dot represents an islet and colors indicate donors. No significant differences were detected by the Mann–Whitney U test. (**D**) Number of genes versus UMIs per capture spot (10 μm) in spatial transcriptomics and Slice-seq data. (**E**) Immunostaining (IH) of a representative islet, with corresponding Ca^2+^ activity quantified in regions of interest (ROIs) at low and high glucose conditions with or without amino acids. (**F**) Representative Ca^2+^ traces for each cell type obtained from the slice in **E**. (**G**) Ca^2+^ event frequency for α (N=237 in 20 islets), β (N=648 in 26 islets), δ (N=59 in 15 islets), acinar (N=2070) and ductal (N=94) cells in 7 ND-Lean donors. Mann–Whitney U test with Bonferroni correction; *p < 0.05, **p < 0.01. (**H**) Gene expression in Slice-seq data from three islets within the same tissue slice. From left to right: islet cell type composition, β cell activity (10 mM glucose), and β cell pseudobulk expression (within the islet) of selected regulators of β cell excitability, secretory machinery, and paracrine signaling. The size of these islets is comparable to that of small endocrine objects recently described in ^11^.

To assess the robustness of this approach, we compared samples processed with Slice-seq or spatial transcriptomics alone. Transcriptomic quality metrics were comparable between methods, with no significant differences in detected genes, UMI counts, or spatial capture coverage, and no evidence of reduced tissue coverage or loss of capture beads (**Fig. 5C-D**; **Supp. Fig. S5B**). Together, these results indicate that the Slice-seq workflow, including acute slice preparation and live cell Ca^2+^ imaging, does not compromise transcriptomic data quality.

Cytosolic Ca^2+^ dynamics were recorded under three perfusion conditions to stimulate islet-cell activity: low glucose, high glucose, and high glucose supplemented with amino acids (**Supp. Video 1**). Regions of enhanced activity corresponding to islets could be identified based on their glucose-responsive behavior (**Fig. 5E**), and by their overlap with sometimes visible peri-islet structures. Regions of interest (ROIs) were defined as described (see **Methods**) ^48^ and classified into cell types based on Ca^2+^ dynamics and morphology. ROI assignments were validated by immunohistochemistry: hormone-positive regions overlapped with endocrine-like Ca^2+^ dynamics, whereas hormone-negative regions lacked glucose-stimulated activity.

We obtained representative traces for all major pancreatic cell types, including islet β, α, and δ cells (**Fig. 5F**, yellow, magenta, gray traces), as well as acinar and ductal cells (**Fig. 5F**, blue, cyan) which have remained comparatively understudied in human tissue. For each trace, we extracted parameters representing Ca^2+^ event frequency, half-width, and event size (**Supp. Fig. S5C**). Overall, endocrine cells were characterized by frequent Ca^2+^ spikes that responded to changes in glucose conditions, whereas acinar and ductal cells displayed broader and slower oscillatory dynamics largely insensitive to glucose conditions.

Quantification of Ca^2+^ event frequency confirmed cell type-specific responses to metabolic stimulation (**Fig. 5G**; **Supp. Fig. S5D**). β cells showed increased event frequency under high glucose, further enhanced by amino acid supplementation. In contrast, α cells were most active under low glucose and suppressed by glucose stimulation, whereas δ cells remained active throughout the protocol without significant changes in event frequency. Similarly, ductal and acinar cells did not show significant changes in event frequency across conditions. Endocrine cells exhibited short Ca^2+^ median half-widths (α: 5.1 s [IQR 3.5-12.0]; β: 5.2 s [IQR 3.5-8.4]; δ: 4.7 s, [IQR 2.6-6.5]), while exocrine cells had significantly longer events (acinar: 10.3 s [IQR 9.0-12.3]; ductal: 17.4 s [IQR 12.6-23.5]) (**Supp. Fig. S5D**), in line with previous observations in rodent experiments ^49–51^. Together, these results provide a broad survey of Ca^2+^ dynamics across all major human pancreatic cell types.

Islet activity was variable across and within acute slice experiments, with some islets showing higher Ca^2+^ activity and larger fraction of active cells than others. Slice-seq is particularly suited to investigate this heterogeneity, especially in small endocrine structures missed by classical studies using isolated islets. Representative islets from the same Slice-seq sample illustrate that the method can resolve both compositional and transcriptional differences in small islets (∼50 μm effective diameter; **Fig. 5H**), while larger canonical islets are shown in **Supp. Fig. S5E**. Heterogeneous expression of Ca^2+^ regulators, incretin receptors, and transcription factors was observed in β cells from islets with different levels of β cell activity (**Fig. 5H**; **Supp. Fig. S5E**). Among these features, islets enriched in α cells tended to show higher fractions of active β cells together with increased expression of incretin receptors in β cells. Although exploratory, these observations highlight that islets with different composition may have different functional properties, and support the relevance of paracrine regulation of β cell activity in human islets. Thus, together with improving the functional characterization of endocrine and exocrine cell types in the human pancreas, our studies provide evidence of heterogeneity in Ca^2+^ activity across islets that is associated with differences in endocrine cell composition and islet-specific gene expression programs.

### Extra-islet β cells exhibit altered metabolic coupling

To extend these observations beyond islet-level heterogeneity, we next used Slice-seq to investigate β cells located outside canonical islets (“extra-islet β cells”). Although small β cell clusters and extra-islet β cells may comprise a significant fraction of β cell mass in humans, their physiological relevance remains unresolved. Hence, we investigated if Slice-seq can recover functional and transcriptomic profiles of extra-islet β cells. To this end, we identified extra-islet β cells in Slice-seq experiments (**Fig. 6A**) and assessed their activity using Ca^2+^ imaging (**Fig. 6B**). To facilitate direct comparisons, we analyzed cells within the same tissue slices, and identified islet and extra-islet β cells based on their Ca²⁺ traces, followed by post hoc validation with immunostaining. Islet β cells showed significant glucose responsiveness as measured by increases in Ca^2+^ event frequency, which was further enhanced by amino acids. In contrast, extra-islet β cells showed blunted glucose responses that were only partially restored by amino acid supplementation (**Fig. 6C**; **Supp. Fig. S6A**)

**Figure 6.**
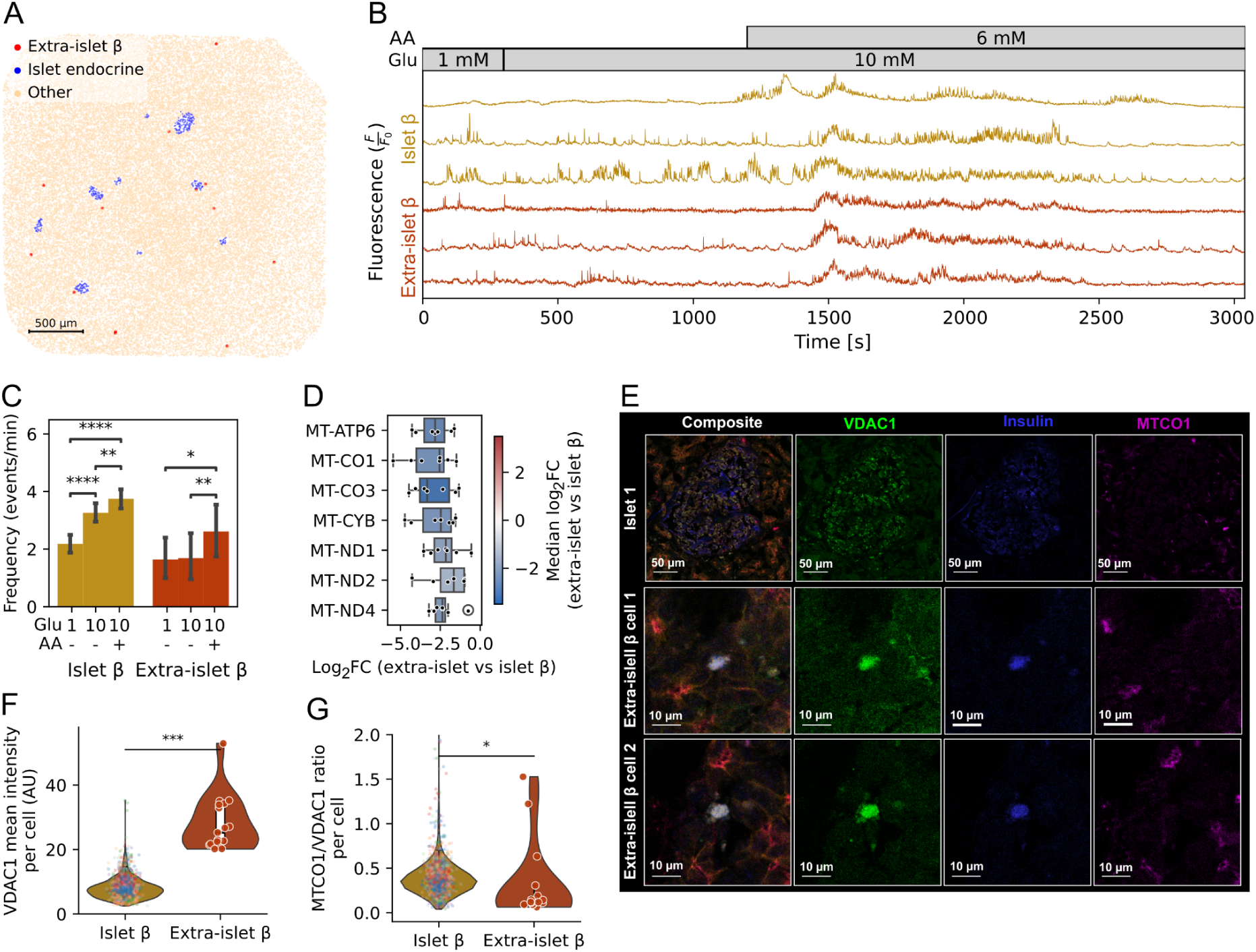
Profiling of extra-islet β cells. (**A**) Distribution of β cells in spatial transcriptomics. (**B**) Representative Ca^2+^ traces of islets and extra-islet β cells from the same slice. (**C**) Frequency of Ca^2+^ firing events in islet and extra-islet β cells (N=45 extra-islet cells, N=648 islet cells, 7 donors). *p < 0.05, **p < 0.01 (Mann–Whitney U test with Bonferroni correction). (**D**) Top differentially expressed genes between extra-islet and islet β cells based on spatial transcriptomics from ND-Lean donors. Each dot represents a donor average. Analysis was restricted to samples containing ≥10 β cells per group. (**E**) Immunostaining images of two representative extra-islet β cells and one islet. (**F**) Quantification of mitochondrial mass (VDAC1) in segmented β cells. (**G**) Quantification of complex IV (MTCO1) to VDAC1 in segmented β cells. In F-G dot colour indicates cells from independent islets.

To investigate molecular factors underlying these functional differences, we analyzed spatial transcriptomics profiles from the Slice-seq experiments. Extra-islet β cells showed reduced expression of mitochondrial-encoded genes involved in oxidative phosphorylation, specifically complex I (MT-ND1/4) and complex IV (MT-CO1/3) **(Supp. Fig. S6B)**. We confirmed reduced expression of these genes in extra-islet β cells in an expanded dataset including all ND-Lean donors profiled by spatial transcriptomics (**Fig. 6D**), pointing to potential altered metabolic coupling in these cells.

To validate these findings, we performed immunofluorescence staining for mitochondrial mass (VDAC1) and complex IV (MT-CO1), and segmented islet and extra-islet β cells based on insulin and E-cadherin expression (**Fig. 6E**; **Supp. Fig. S7**; see **Methods**). We found that extra-islet β cells exhibited increased mitochondrial mass (∼3.2-fold, P<0.001), but reduced MTCO1/VDAC1 ratio (P=0.019), indicating decreased complex IV abundance relative to mitochondrial content (**Fig. 6F-G**). Together, these findings suggest that β cell metabolic state is influenced by tissue context, with extra-islet β cells exhibiting a decoupling between mitochondrial mass and oxidative capacity that may underlie the reduced glucose responsiveness observed in Ca^2+^ imaging.

## DISCUSSION

The pancreas is a highly organized secretory organ in which endocrine, exocrine, stromal, and immune cells interact closely to regulate tissue function and homeostasis. Single-cell studies have revealed extensive regulatory and transcriptional heterogeneity across pancreatic cell types ^29,52^, but linking these cellular states to physiological activity *in situ* has remained challenging. Here, we developed Slice-seq, a multimodal approach that links Ca^2+^ activity to spatial transcriptomics in acute human pancreatic tissue slices. By preserving tissue context, Slice-seq enabled functional characterization of all major human endocrine and exocrine cell types, revealed heterogeneity in islet activity and gene expression as well as defective glucose responsiveness and mitochondrial oxidative capacity in extra-islet β cells. In parallel, integration of single-nuclei and spatial transcriptomics uncovered exocrine remodeling in obesity, partially overlapping with changes observed in T2D.

Compared with islets, exocrine cells have remained substantially less studied in metabolic disease, in part due to technical challenges in profiling acinar-rich tissue and the lack of stable *in vitro* models ^53^. Hence, studies of human tissue remain crucial for understanding exocrine alterations in disease ^37^. In contrast to the compensatory increase in β cell secretory function during early diabetes adaptation ^56^, our data indicates that obesity and T2D are associated with downregulation of mature functional programs in acinar and ductal cells together with induction of inflammatory and stress-response transcripts. These changes are not accompanied by major shifts in cell-type composition or macrophage abundance and polarization, suggesting that systemic metabolic stressors, including hyperglycemia and dyslipidemia ^56^, or local microenvironmental cues such as islet-derived secreted factors ^57^, may contribute to exocrine remodeling beyond immune infiltration. Our findings also resonate with previous mouse studies showing that disruption of transcriptional programs maintaining acinar differentiation can generate a pro-inflammatory state, even in the absence of increased immune-cell infiltration ^58^. The similarities with the remodeling observed here support the possibility that related epithelial-intrinsic processes contribute to exocrine alterations in obesity and T2D. Although acinar and ductal subtypes displayed broadly consistent responses, REG+ acinar cells ^29^ exhibited a comparatively milder phenotype, suggesting that distinct acinar cell states may differ in their response to metabolic stress. Understanding the mechanisms regulating these states may therefore provide new avenues for targeting inflammatory responses in exocrine pancreatic disease.

Recent volumetric endocrine mass assessments using 3D imaging ^10^, together with high-throughput histological analyses ^11^, have shown that almost 50% of endocrine objects do not conform to stereotypical islet structures, extending long-standing observations of heterogeneity in human islet architecture and composition ^43,59^. Such structural diversity may contribute to molecular and functional heterogeneity across islets ^60^. However, connecting cytoarchitecture to transcriptional states and physiological activity has remained challenging in single-cell genomic studies, due to the need for islet and tissue dissociation. Our spatial transcriptomics analyses recapitulated these seminal findings while further linking β cell-rich islets to increased expression of regulators of β cell excitability. Leveraging the rich molecular and spatial information captured by spatial transcriptomics, we further profiled islet gene expression in donors with T2D. In δ cells, we observed an upregulation of actin cytoskeleton remodeling pathways in the context of local macrophage infiltration. Given that δ cells form filopodia-like contacts with neighboring α and β cells ^38^, these changes could impact paracrine β cell regulation in T2D. Enhanced actin cytoskeleton signaling has recently been reported in proteomic studies of human T2D islets ^61^, while δ cell composition has also been associated with T2D genetic risk in large-scale islet imaging studies ^14^. Together, these findings suggest that endocrine cells can express context-dependent transcriptional programs and support the idea that local cell interactions within islets regulate endocrine cell states and activity.

Slice-seq enabled functional characterization of islet heterogeneity in its tissue context, including small endocrine objects difficult to capture in isolated islet experiments. The association between α cell abundance, incretin signaling pathways, and increased β cell activity recapitulates known features of paracrine α-to-β-cell communication in islets ^62,63^, while linking these interactions to local transcriptional programs. Previous Patch-seq studies identified β cell states with distinct electrophysiological and transcriptional profiles in dissociated human islet cells ^63^, but whether these states are uniformly distributed across islets remained unclear. Our observations suggest that part of this heterogeneity is linked to islet composition and local microenvironment, further connecting islet architecture to β cell functional and transcriptional heterogeneity across human islets.

This heterogeneity further extended to extra-islet β cells, which showed blunted glucose responsiveness together with reduced mitochondrial oxidative metabolism. Extra-islet β cells lack the physical and paracrine interactions with neighboring endocrine cells characteristic of canonical islets while also being exposed to the exocrine microenvironment ^60^. Given that β cells are susceptible to the accumulation of oxidative stress-associated signatures with age ^64^, the reduced oxidative capacity observed here suggests that below a certain size, small endocrine objects may become increasingly vulnerable to metabolic stress. Recent studies have linked extra-islet β cells to T1D pathophysiology ^11,65^, yet little is known about their function in human tissue. Future experiments using juvenile tissue may help clarify whether the features identified here are intrinsic to extra-islet β cells or progressively acquired during metabolic stress and aging.

Slice-seq is performed in acute *ex vivo* tissue slices, which preserve physiological interactions more effectively than isolated cells, but do not fully recapitulate *in vivo* physiology. In addition, the short ischemic times required for acute slice studies limit sample size, and larger cohorts will be needed to map exocrine cell states across metabolic and disease conditions, ideally under exocrine stimulatory conditions. Unlike dissociated-cell approaches such as Patch-Seq, Slice-seq relies on integration across imaging, immunohistochemistry, and spatial transcriptomics, making direct mapping at single-cell resolution challenging. Nevertheless, preservation of local tissue organization across adjacent sections enables characterization of cell-type-associated functional states within the same islets and spatial regions. Finally, integration of Slice-seq with long-term slice culture, perturbation experiments, molecular recording, or repeated sampling may further resolve dynamic transitions in pancreatic cell states.

In conclusion, our study comprehensively maps endocrine and exocrine cell states in health and metabolic disease and establishes a framework for interrogating human pancreatic biology *in situ* by linking transcriptional states to dynamic physiological activity in acute tissue slices.

## METHODS

### Sample procurement

Human pancreatic samples used in this study were collected across 4 different academic centers (42 donors in total, 9 from Uppsala University in Sweden, 12 from the ADI IsletCore in Canada, 14 from the Medical University of Vienna in Austria and 7 from the University of Pisa in Italy). All studies have been approved by local IRBs, and following SOPs and study protocols detailed below. Donor characteristics (age, sex, BMI, HbA1c) and available clinical information for each donor are listed in **Supp. Table 1A**. The mean age, BMI and HbA1c, as well as number of donors by sex in each study group, are summarized in **Supp. Table 1B**. Classification of donors as nondiabetic, pre-T2D or T2D was based on medical record information or postmortem HbA1c value. Donors with prior T2D diagnosis per medical record or HbA1c≥6.5 were classified as T2D, donors without prior T2D diagnosis and 5.7≤HbA1c≤6.4 were classified as pre-T2D, and donors without prior T2D diagnosis and HbA1c≤5.6 (or HbA1c unavailable) were classified as nondiabetic. Non-diabetic donors were further classified as non-diabetic lean (BMI<30) or non-diabetic obese (BMI≥30).

This study was approved by the Swedish Ethics Review Authority (2025-08902-01, Dnr 2023-01845-01, Dnr 2015/444). A complete de-identified demographics table is provided in **Supp. Table 1**. Fresh human pancreatic tissue as well as snap frozen pancreatic tissue from donors with and without diabetes were obtained from Uppsala University through the group of Olle Korsgren.

Alberta Diabetes Institute IsletCore at the University of Alberta in Edmonton (www.isletcore.ca) samples were procured with the assistance of the Alberta Health Services Give Life Alberta program, the Ontario Health Trillium Gift of Life Network (TGLN), Quebec Transplant, and other Canadian organ procurement organisations. The Human Research Ethics Board at the University of Alberta approved tissue collection (Pro00013094). All donors’ families gave informed consent for the use of pancreatic tissue in research.

Pancreatic tissue was obtained from deceased organ donors at the Division of Transplantation, Medical University of Vienna, in accordance with institutional standard operating procedures. Organ donation in Austria operates under a presumed consent (opt-out) framework, and no donor included in this study had a documented objection to organ donation. The retrieval and use of tissue samples was approved by the institutional review board at the Medical University of Vienna (EK# 2191/2016). Fresh pancreatic tissue specimens were collected during organ procurement from donors whose pancreas was not allocated for transplantation. All organs were perfused with histidine-tryptophan-ketoglutarate (HTK) solution as part of routine retrieval, preserving cellular integrity and inhibiting autolytic processes.

Human OCT-embedded pancreatic tissue samples from donors with and without T2D were provided by Piero Marchetti (University of Pisa, Italy).

### Study design overview

Human pancreatic tissue was processed using two complementary workflows. Fresh-frozen tissue was used for snRNA-seq, immunostaining and spatial transcriptomics, enabling characterization of cell types, transcriptional states and their spatial organization. In parallel, a subset of samples was processed into viable acute tissue slices for live cell Ca^2+^ imaging, after which the imaged slice was subjected to spatial transcriptomics and immunostaining (Slice-seq), linking functional activity to spatial gene expression.

### Single-nuclei RNA sequencing

Isolation of pancreatic single nuclei was performed according to a previously published protocol with some modifications ^29^. Fresh frozen pancreatic tissue was sectioned using a cryostat (CM3050S, Leica, Germany) into 5 x 5 x 0.3 mm^3^ pieces, handled with precooled forceps, and stored on dry ice until further processing. The tissue was placed in a 2 ml glass tissue grinder (PSTL LC, Kimble Chase, New Jersey, USA) and homogenized with one stroke of “loose” pestle in 1 ml of citric acid-based buffer (Sucrose 0.25 M, Citric Acid 25 mM, Hoechst 33342 1 µg/ml, RiboLock RNase inhibitor 800 mU/µl, Baseline-ZERO^TM^ DNase 5 mU/µl). The tissue was incubated on ice for 5 min, followed by an additional 5 strokes and another 5 min incubation on ice.

The final homogenization step consisted of 5 strokes of a “loose” pestle followed by 5 strokes of a “tight” pestle. The homogenate was filtered through a 30 µm cell strainer (pluriselect, Germany) and centrifuged at 500 g for 6 min at 4 °C (Avanti J-15R, Backman Coulter, California, USA). The supernatant was carefully removed, and the nuclei were washed in 1 ml of citric acid-based buffer, followed by another centrifugation. The supernatant was removed, and the nuclei were resuspended in 300 µl of citric acid-based buffer. An equal volume of high-density buffer (Sucrose 1.5 M, Citric Acid 250 mM, RiboLock RNase inhibitor 470 mU/µl, Baseline-ZERO^TM^ DNase 2.9 mU/µl) was slowly added to the bottom of the tube. The sample was centrifuged at 10,000 g for 11 min at 4 °C. After removing the supernatant, the nuclei were resuspended in 300 µl of citric acid-based buffer and filtered through a 30 µm cell strainer. The nuclei were further purified using the nuclei enrichment protocol Anti-Nucleus MicroBeads, according to the manufacturer’s instructions (Miltenyi Biotec, 130-132-997). To assess the quality and purity of isolated nuclei, they were imaged using a ZOE Fluorescent Cell Imager (Bio-Rad, California, USA) with a 20x objective.

Single nuclei samples were fixed following the Evercode^TM^ Nuclei Fixation v3 protocol (ECFN3300, Parse Bioscience). The nuclei were loaded according to the sample loading table provided by Parse Bioscience. A total of 32 samples were loaded with an expected yield of 100,000 single nuclei sequenced. Library preparation was performed using Evercode^TM^ WT v3 with Integra Assist Plus instructions (ECWT3300, Parse Bioscience). Final libraries were quantified using a Qubit fluorimeter (Invitrogen, California, USA) and the size distribution was checked using a Tapestation system (High Sensitivity D5000 assay, Agilent, 5067). All centrifugation steps during fixation and library preparation were made at 500 × g for 6 min at 4 °C (Avanti J-15R, Beckman Coulter, California, USA).

### Spatial transcriptomics

Spatial transcriptomics experiments were performed according to the Curio Seeker 3 x 3 Spatial Mapping Kit v2.2 (Curio Bioscience, California, USA) ^66^.

Fresh frozen pancreatic tissue was cryo-sectioned to a thickness of 10 µm slides using a cryostat (CM3050S, Leica, Germany) set at a chamber temperature of -26 °C and a sample holder temperature of -20 °C. Sections were then transferred to a Curio Tile using an in-house Tissue Stamper. The Tissue Stamper is composed of ThorLabs components (ThorLabs, New Jersey, USA), a 4K WiFi Digital Microscope (Canada, California, USA) and self-3D-printed components. The crucial component of the stamper is a thin layer of polydimethylsiloxane (PDMS), which attracts optimal cutting temperature (OCT) and/or tissue due to its hydrophobic properties, allowing for electrostatic interactions (surface charge accumulation). After applying the tissue to the Curio Tile, tissue was melted onto the tile. The tile with melted tissue was transferred to 200 µl of hybridisation buffer and incubated for 30 min at room temperature (RT). Following incubation, the tile was washed in reverse transcription wash buffer and transferred to 200 µl of reverse transcription reaction mix and incubated first for 10 min at RT and then for 30 min at 52 °C. Tissue digestion was performed by the addition of 200 µl of tissue clearing reaction mix and incubating it for 30 min at RT. After digestion, the beads were washed twice with a bead wash buffer and pelleted at 3000 g for 2 min.

Second-stranded synthesis was carried out by resuspending the beads in 200 µl of second-strand mix and incubating for 1 h at 37 °C. The beads were washed twice with a bead wash buffer and resuspended in the cDNA amplification mix. The reaction was split into 50 µl aliquots and amplified in the thermocycler using the following protocol: 98 °C for 2 min; four cycles of 98 °C for 20 sec, 65 °C for 45 sec, 72 °C for 3 min; 9-13 cycles of 98 °C for 20 sec, 67 °C for 20 sec, 72 °C for 3 min; final 72 °C for 5 min and 4 °C hold. The amplified cDNA was purified twice with SPRIselect beads (Beckman Coulter, B23318) at a 0.7x beads-to-sample ratio, eluting in a final volume of 20 µl. Quantitative and qualitative assessment of cDNA was performed using High Sensitivity D5000 TapeStation assay (Agilent, 5067).

For Illumina sequencing library preparation, 700 pg of cDNA was used. The Nextera^®^ XT kit (15032350, Illumina, California, USA) was used for tagmentation, followed by amplification with barcoded index primers. Libraries were purified twice with SPRIselect beads (B23318, Beckman Coulter, California, USA) at a 0.8x beads-to-sample ratio, eluting in a final volume of 20 µl.

### Immunostaining

The subsequent slide of spatial transcriptomics was used for an immunohistochemistry assay. Pancreatic tissue slides were sectioned at a thickness of 10 µm and fixed in 4% PFA for 15 min at RT. Slides were then washed three times in PBS and permeabilised/blocked with a solution containing 0.3% Triton X-100, 5% BSA in PBS for 30 min at RT.

Tissue slides were incubated overnight at 4 °C with primary antibodies: rat anti-insulin (1:50; R&D Systems, MAB1417), mouse anti-glucagon (1:200; Sigma-Aldrich, G2654), and rabbit anti-somatostatin (1:100; Sigma-Aldrich, SAB4502861). Following incubation, the tissue slides were washed three times with PBS and incubated with secondary antibodies for 2 h at RT: goat anti-rat Alexa Flour^TM^ 488 (1:500; Invitrogen, A48262), goat anti-rabbit Alexa Flour^TM^ 568 (1:500; Invitrogen, A11036) and goat anti-mouse Alexa Flour^TM^ 647 (1:500; Invitrogen, A32728). Tissue slides were washed three times with PBS, and to assess morphology, they were stained with DAPI (0.1 µg/mL in PBS; Merck, D9542). Finally, the slides were mounted with ProLong™ Diamond Antifade mounting medium (ThermoFisher, P36965). Slides were imaged on the CellDiscoverer 7 (Zeiss, Germany) using a Plan-Apochromat 20x/0.7 objective equipped with an Axiocam 712 mono camera in Tile mode.

### Preparation of viable tissue slices

Pancreatic tissue was acquired as specified, with one part of it directly frozen for the snRNA-seq, and the other processed into viable acute human pancreatic tissue slices, as described previously ^26,66,67^.The human tissue was sectioned into small cubes (∼5 mm^3^), embedded in 3.8% low-melting-point agarose (Lonza, 11540737) dissolved in extracellular solution (ECS), consisting of the following components (in mM): NaCl 125, NaHCO₃ 26, glucose 4, lactic acid 6, myo-inositol 3, KCl 4, Na-pyruvate 2, CaCl₂ 2, NaH₂PO₄ 1.25, MgCl₂ 1, and ascorbic acid 0.25, at 40 °C. Immediately after embedding, the agarose with tissue was cooled on ice and sliced into 140-µm-thick sections using a vibratome (VT 1000 S, Leica Microsystems, Germany) in ice-cold ECS supplemented with Aprotinin Protease Inhibitor (10 µg/ml; Thermo Scientific™, 78432) and RiboLock RNase Inhibitor (0.01 U/µl; Thermo Scientific™, EO0381).

The tissue slices were then loaded with the Ca^2+^ indicator Calbryte 520-AM (6 µM; AAT Bioquest, 20651) for 1 hour in HEPES-buffered ECS (HB-ECS). HB-ECS consisted of (in mM): NaCl 125, NaHCO₃ 10, HEPES 10, glucose 4, lactic acid 6, myo-inositol 3, KCl 4, Na-pyruvate 2, CaCl₂ 2, NaH₂PO₄ 1.25, MgCl₂ 1, and ascorbic acid 0.25, titrated to pH 7.4 using 1 M NaOH. This solution was supplemented with 0.03% Pluronic F-127 (w/v), Aprotinin Protease Inhibitor (10 µg/ml; Thermo Scientific™, 78432), RiboLock RNase Inhibitor (0.01 U/µl; Thermo Scientific™, EO0381), and 0.12% dimethyl sulfoxide (DMSO) at RT in the dark. The slices were incubated on a rotor-shaker at a constant speed of 100 rpm.

### Cytosolic Ca^2+^ imaging of acute slices

The individual pancreatic tissue slice was transferred into an imaging chamber equipped with a super-perfusion system. Slices were imaged on confocal microscope Zeiss 700 (galvo scanner, 3 Hz, 256 x 256, objective: PL APO 20x/0.8 air) or NikonA1R (resonant scanner, 15 Hz, 512 x 512, objective: APO LWD 20x/0.95 water), accounting for the pixel size to be around 1 µm ^26^. Calbryte 520 was excited by a 488 nm argon laser, and the emitted light was collected with GaAsP PMT detectors in the 500-700 nm range. We imaged a layer of cells approximately 30 µm deep in the tissue. During imaging, the tissue was constantly super-perfused with ECS containing 1 or 10 mM glucose with or without the addition of an amino acid cocktail (glutamate, arginine and alanine, 2 mM each) at 37 °C.

### Slice-seq methodology

Following live Ca²⁺ imaging, acute pancreatic tissue slices were re-embedded in OCT while preserving their orientation for subsequent cryo-sectioning. For cryo-sectioning, the cryostat (CM3050S, Leica, Germany) was set to a chamber temperature of -26 °C and a sample holder temperature of -20 °C. Successful sectioning of a 140 µm acute tissue slice requires proper alignment of the cutting blade and the tissue section. Excess OCT was removed in 50–100 µm increments until the tissue became visible, after which sectioning proceeded at 10 µm intervals. Cut slices were examined under a stereomicroscope to verify the presence and integrity of the tissue. Consecutive sections were collected throughout the tissue for future use, with the 9th-10th sections used for spatial transcriptomics, as these correspond to the imaging plane. Remaining 10 µm sections were collected until exiting the tissue slice to verify tissue depth and maximize tissue utilization.

### Immunostaining of VDAC1 and MTCO1 in human pancreatic tissue slices

Cryosections (8 µm) were mounted onto charged slides and fixed in ice-cold 4% PFA in PBS for 10 minutes at room temperature, quenched with 0.1 M glycine in PBS, and permeabilised with 0.05% Triton X-100 in PBS. Sections were blocked with 10% normal goat serum (Vector Laboratories, SP-2001) + 1% BSA in PBS for 1 hour at room temperature. Primary antibodies: MTCO1 (Abcam, ab14705, 1:200), VDAC1 (Abcam, ab14734, 1:100), insulin (DSHB, GN-ID4, 1:100), and E-cadherin (Cell Signalling Technology, 24E10, 1:500); were incubated overnight at 4 °C. The following Alexa Fluor-conjugated secondary antibodies (all 1:500) were applied for 1 hour at room temperature: goat anti-rat Alexa Fluor^TM^ 405 (Fisher, 17133711), goat anti-mouse IgG2b Alexa Fluor^TM^ 488 (Thermo Fisher Scientific, A21141), goat anti-rabbit Alexa Fluor^TM^ 568 (Thermo Fisher Scientific, A11036), and goat anti-mouse IgG2a Alexa Fluor^TM^ 647 (Thermo Fisher Scientific, A21241). Sections were mounted with ProLong Diamond Antifade Mountant (Thermo Fisher Scientific, P36961). No-primary and subclass specificity controls were included. Images were acquired on a Zeiss LSM980 confocal microscope (Plan Apo 20x/0.8 NA; 0.1235 µm/pixel). Whole-tissue single-cell segmentation and identification of islet and extra-islet β cells were performed using a custom Cellpose-based image analysis pipeline. Mitochondrial content was quantified as the mean MTCO1/VDAC1 fluorescence intensity ratio per cell. Statistical differences between islet and extra-islet β cells were assessed using a linear mixed effects model with cell type as a fixed effect and islet identity as a random effect, applied to log-transformed values, with Benjamini-Hochberg FDR correction across three markers.

### Processing and analysis of Ca^2+^ imaging data

Ca^2+^ imaging data were processed using the previously described automated analysis pipeline ^26^ with key steps summarised here. Briefly, recordings were handled as three-dimensional (T x X x Y) arrays and, where necessary, corrected for motion using template-based frame alignment implemented as in CaImAn ^68^ after temporal rebinning to reduce noise. Regions of interest (ROIs) corresponding to individual cells were identified using a semi-automated procedure that applied bandpass filtering to a representative image (combining mean and high-percentile projections) to enhance local intensity variations at the spatial scale of islet cells (∼10 µm). Pixels were grouped into ROIs by assigning them to local intensity maxima. This approach minimises manual intervention while enabling robust detection of large cell populations within high-volume recordings.

Fluorescence traces were analysed under the assumption of photon-counting (Poisson-like) noise and transformed into z-scores to normalize signal fluctuations. Slow baseline components were estimated using cascaded second-order section (SOS) filtering implemented in scipy.signal module ^68^, excluding outliers at each step to reduce distortion. ROI signals were computed as the sum of pixel intensities to preserve correct noise scaling. Event detection was performed across multiple temporal scales using sequential filtering, with candidate events identified by thresholding (z > 4) and subsequently refined through a multi-scale consistency criterion (“event distilling”). Events were defined by their onset time, amplitude, and half-width, and only those reproducibly detected across timescales and exceeding duration and boundary criteria were retained, reducing false positives while preserving sensitivity to dynamics spanning milliseconds to minutes.

### Immunostaining analysis

Multi-channel fluorescence microscopy images (.tif format) were processed using a custom Python pipeline (Python 3.11.13) with a tiff file for image reading. Individual channels (insulin/β, somatostatin/δ, glucagon/α, and DAPI/nuclei) were normalized using percentile-based contrast enhancement (90th-99th percentiles) and merged into RGB images with channel-specific color mapping. Cell and nuclei segmentation was performed using Cellpose 4.0.6 ^69^ using GPU acceleration. Segmentation parameters were optimized for pancreatic tissue: flow threshold = 10, cell probability threshold = -3.0, tile normalization block size = 500 pixels, and 5000 iterations. Nuclei-specific segmentation used flow threshold = 0.4 and cell probability threshold = 0. Segmentation masks were filtered to remove artifacts smaller than 200 pixels and validated against reference nuclei positions using spatial overlap criteria (5-pixel tolerance). Cell classification was performed by quantifying cytoplasmic signal intensity (excluding DAPI-thresholded nuclear regions dilated by 1 pixel) using Otsu thresholding per channel. Cells were classified as α (glucagon-positive), β (insulin-positive), or δ (somatostatin-positive) if ≥ 5% of cytoplasmic pixels exceeded the channel-specific threshold. Per-cell measurements included total cell area, nuclear area, nuclear-to-cell area ratio, mean signal intensity per channel, fraction of positive pixels, and spatial coordinates. Quantitative analysis and visualization were performed using scikit-image (v0.25.2), pandas (v2.3.3), and matplotlib (v3.10.6). Cells with areas outside physiological ranges or insufficient cytoplasmic pixels (<10 pixels) were excluded from analysis.

### Processing and normalization of snRNA-Seq data

Parse Biosciences libraries were reprocessed with split-pipe pipeline (v1.6.2) with a custom hg38 reference (GRCh38 primary assembly, Ensembl v109). The pipeline performs barcode correction, alignment, deduplication, and transcript quantification, generating gene-by-nucleus count matrices and QC metrics. Doublets were filtered using Scrublet (score < mean + 1 SD). Cells with > 3% mitochondrial transcripts, nuclei with < 300 genes, and genes detected in < 3 nuclei were excluded. Additional doublets were removed based on marker gene co-expression during clustering and annotation. Analyses were performed using Scanpy and AnnData ^70^. Counts per cell were normalized to a total of 10^6^ counts and log-transformed.

### Clustering and cell type annotation of snRNA-seq data

After extensive barcode quality control and filtering, 56,680 nuclei transcriptomic profiles were retained. Cell types and subtypes were annotated using Leiden clustering. We identified 22 major clusters embedded in the UMAP space (**Fig. 1C**), which were annotated based on known marker gene expression (**Supp. Fig. S1A**), including endocrine islet cells (α, β, δ, γ), exocrine cells (acinar and ductal), immune cells (macrophages, plasmablasts, T cells, mast cells), as well as endothelial, stellate, and Schwann cells. Pseudo-bulk expression profiles for each cell type were created by aggregating counts across cells and normalized to counts-per-million (CPM) (**Supp. Table 2**). On average, 17306 genes were expressed (CPM > 1) per cell type. Differential expression analysis identified cell type- and subtype-specific genes (FDR < 0.10) (**Supp. Table 3**).

### Associations between snRNA-seq data and clinical parameters

The proportion of each cell type relative to all nuclei, and of each cell subtype within its primary cell type, was computed per sample. Differences in proportions across groups (ND-Lean, ND-Obese, pre-T2D, and T2D) were assessed using the Mann-Whitney U test. Samples with abnormally low cell counts were excluded (< 50 for acinar, and ductal cells; < 10 for stellate, immune and endothelial cells). Associations between cell subtype proportions and clinical parameters (age, body mass index [BMI], and HbA1c) were assessed using Spearman’s correlation. For transcriptomic analysis, log-transformed pseudo-bulk expression was computed per donor for each cell type. Associations between gene expression and BMI across non-diabetic donors were assessed using Pearson’s correlation to retain information about the magnitude of BMI differences across donors.

### Differential gene expression and pathway enrichment analysis

Differential expression analysis of the snRNA-seq data was performed using both DESeq and memento ^33^. For Memento, sex was included as a covariate. For DESeq2, pseudo-bulk count matrices were generated for each cell type by aggregating counts across all barcodes per donor and gene. Multiple testing correction was performed using the Benjamini–Hochberg method. Gene set enrichment analysis (GSEA) was performed to identify disease-associated pathways, using GO and Reactome as reference databases. Pathways were considered significant at FDR < 0.1 and required at least two contributing genes.

### Trajectory analysis for endothelial cells

Trajectory inference and branching analysis were performed using the Scanpy function dpt, and RNA velocity–based trajectories were computed with scVelo ^70^. Vessel size scoring was derived from the top 50 positively and negatively correlated genes with pseudotime (Spearman’s correlation) along the capillary-to-arterial and capillary-to-venous pseudotime trajectories, using capillary endothelial cells as the root. Scores for positively and negatively correlated genes were calculated separately, and a combined vessel size score was obtained by subtracting the negative score from the positive score. Vessel size categories were assigned based on score percentiles and grouped into ‘large’, ‘medium’, and ‘small’.

### Processing and normalization of spatial transcriptomics data

Spatial transcriptomic sequencing data were processed using the Curio Seeker bioinformatics pipeline. Raw FASTQ files were aligned, spatial barcodes were decoded, and gene expression matrices with corresponding spatial coordinates were generated following the manufacturer’s workflow. The resulting data was exported in H5AD format for downstream analysis with Scanpy and Squidpy ^70,71^. Gene expression, metadata, and spot coordinates were imported for each slide and merged into a single AnnData object. Genes detected in fewer than 10 spots were excluded from further analysis. Counts per spot were normalized to a total of 10^4^ and log-transformed.

### Cell type and subtype deconvolution of spatial transcriptomics data

Cell type and subtype annotation in spatial transcriptomics data was performed using Cell2location ^72^. Cell types were assigned to each spatial spot based on the highest Cell2location abundance estimate. To further resolve non-endocrine subtypes, deconvolution was re-run separately for each sample using reference models built from the corresponding snRNA-seq subtypes. Subtype labels were assigned according to the highest inferred Cell2location abundance estimate and validated by expression of established marker genes. High-confidence islets were identified by combining endocrine marker enrichment and local spatial autocorrelation of Cell2location-derived endocrine abundance estimates. Candidate endocrine spots were grouped into individual islets using spatial clustering and retained according to predefined quality criteria. Endocrine subtype annotation within islets was subsequently refined using Cell2location abundance estimates and local neighbourhood information.

### Cellular neighborhood and enrichment analyses

Cellular neighborhood and enrichment analyses were performed on a per-sample basis using Squidpy ^71^. Spatial neighborhood graphs were constructed using the spatial_neighbors function with 8 nearest neighbors, and cell-type neighborhood enrichment was quantified using the nhood_enrichment function with 500 permutations.

### Association between islet composition and cell type-specific gene expression

To identify genes whose expression varies with islet cellular composition, we assessed the relationship between islet endocrine cell type proportions (e.g., β cell fraction) and gene expression in each cell type. Simulated islets with varying proportions of α and β cells were generated based on snRNA-seq data to define a reference model of composition-driven effects and account for local spatial confounding.

Spearman correlation analysis was performed between log₂-transformed pseudo-bulk gene expression (per cell type per islet) and endocrine cell type proportions (e.g., β cells) in both spatially segmented islets and simulated snRNA-seq islets. Genes exhibiting discordant correlation patterns between spatial and simulated islets were interpreted as being associated with tissue context beyond compositional effects and were selected based on z-scores relative to the null distribution derived from simulated islets.

## DATA AVAILABILITY

Raw sequencing reads have been deposited in NCBI Gene Expression Omnibus (GEO) under accession number (GSEXXXXX) and will be publicly available upon publication.

## CODE AVAILABILITY

The code to reproduce the analyses in this study will be made publicly available in Github together with preprocessed datasets in Zenodo prior to publication.

## ACKNOWLEDGEMENTS

J.C.-S acknowledges support from the Knut and Alice Wallenberg Foundation (Wallenberg Molecular Medicine Fellow), the Swedish Research Council (2021-05109), the Erling Perssons Stiftelse (Swedish Foundations’ Starting Grant) and an EFSD/Novo Nordisk Foundation Future Leaders Award (NNF25SA010613). O. K. acknowledges support of the Swedish Research Council (2023-02221), The Leona M. & Harry B. Helmsley Charitable Trust (2103-05057) and Diabetesfonden (DIA2025-993). P.E.M. holds the Canada Research Chair in Islet Biology. The computational analyses were enabled by resources provided by the National Academic Infrastructure for Supercomputing in Sweden (NAISS), partially funded by the Swedish Research Council through grant agreement no. 2022-06725. We acknowledge the technical support as well as the access to the facilities of Patrik Rorsman’s group at the Department of Physiology, University of Gothenburg. Finally, we are indebted to organ donors and their families for their generous support of scientific research.

## AUTHOR CONTRIBUTIONS

Conceptualization: S.P., Y.X., J.C.-S. Methodology: S.P., Y.X., D.P., M.D., M.S.R., O.K., J.C.-S. Investigation: Y.X., S.P., R.F., A.M., S.I., N.S., M.S., M.G.-T. Validation: S.P., R.F., A.M., K.H. Formal Analysis: Y.X., S.P., J.P., M.G.-T., M.S.R., J.C.-S. Data Curation: Y.X., S.P., D.P., M.D., S.I., N.S., M.S. Visualization: Y.X., S.P., A.M., J.P., J.C.-S. Resources: D.P., S.I., N.S., M.S., M.T., G.G, J.D., P.P.S., L.M., P.M., P.E.M., M.S.R., O.K. Supervision: L.M., P.M., P.E.M., M.S.R., O.K., J.C.-S. Funding Acquisition: L.M., P.M., P.E.M., M.S.R., O.K., J.C.-S. Writing - Original Draft: Y.X., S.P., J.C.-S. Writing - Review & Editing: All authors.

