## Supplementary Materials for "Spatial and functional mapping of the human pancreas reveals endocrine and exocrine cell states in health and metabolic disease"

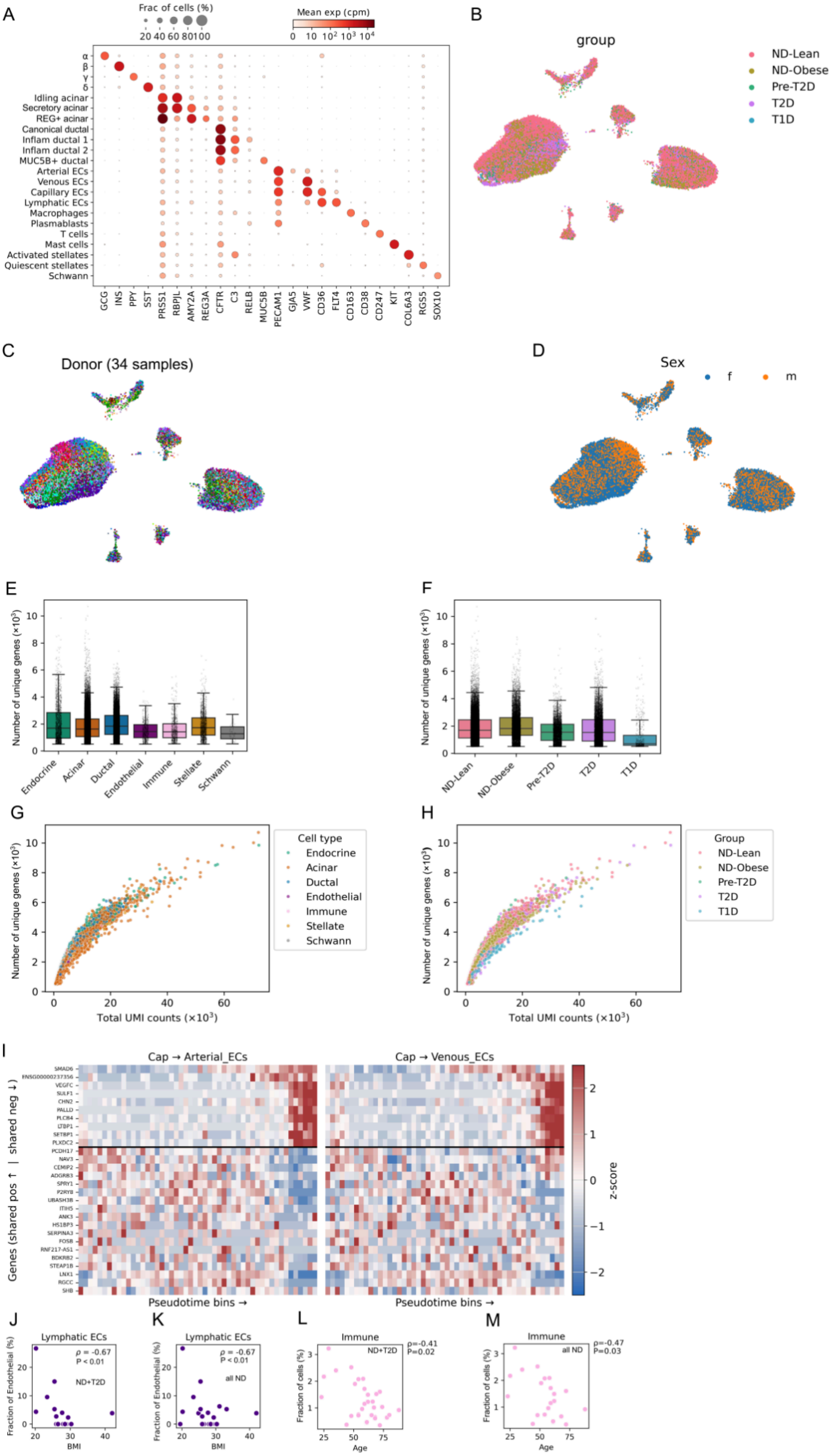
A
Frac of cells (%)
20 40 60 80 100
Mean exp (cpm)
0 10 10² 10³ 10⁴
α
β
γ
δ
Idling acinar
Secretory acinar
REG+ acinar
Canonical ductal
Inflam ductal 1
Inflam ductal 2
MUC5B+ ductal
Arterial ECs
Venous ECs
Capillary ECs
Lymphatic ECs
Macrophages
Plasmablasts
T cells
Mast cells
Activated stellates
Quiescent stellates
Schwann
GCG INS PPY SST PRSS1 RBPJL AMY2A REG3A CFTR C3 RELB MUC5B PECAM1 GJA5 VWF CD36 FLT4 CD163 CD38 CD247 KIT COL6A3 RGS5 SOX10
B
group
ND-Lean
ND-Obese
Pre-T2D
T2D
T1D
C
Donor (34 samples)
D
Sex
f
m
E
Number of unique genes (×10³)
Endocrine
Acinar
Ductal
Endothelial
Immune
Stellate
Schwann
F
Number of unique genes (×10³)
ND-Lean
ND-Obese
Pre-T2D
T2D
T1D
G
Number of unique genes (×10³)
Total UMI counts (×10³)
Cell type
Endocrine
Acinar
Ductal
Endothelial
Immune
Stellate
Schwann
H
Number of unique genes (×10³)
Total UMI counts (×10³)
Group
ND-Lean
ND-Obese
Pre-T2D
T2D
T1D
I
Cap → Arterial_ECs
Cap → Venous_ECs
Genes (shared pos ↑ | shared neg ↓)
Pseudotime bins →
Pseudotime bins →
z-score
J
Lymphatic ECs
ρ = -0.67
P < 0.01
ND+T2D
Fraction of Endothelial (%)
BMI
K
Lymphatic ECs
ρ = -0.67
P < 0.01
all ND
Fraction of Endothelial (%)
BMI
L
Immune
ND+T2D
ρ=-0.41
P=0.02
Fraction of cells (%)
Age
M
Immune
all ND
ρ=-0.47
P=0.03
Fraction of cells (%)
Age

**Supplementary Figure 1. Analysis of snRNA-seq data. (A)** Normalized expression of marker genes for cell types and subtypes. **(B–D)** UMAP plots showing the distribution of nuclei by group (B), donor (C), and sex (D). **(E-F)** Number of unique genes detected across cell types (E) and donor groups (F). **(G-H)** Relationship between the number of detected genes and total UMI counts per nucleus across cell types (G) and donor groups (H). **(I)** Heatmap of vessel size scores based on the top 50 genes positively and negatively correlated with the capillary-to-arterial (left) and capillary-to-venous (right) pseudotime trajectories. **(J-K)** Fraction of lymphatic ECs relative to total ECs versus BMI in all donors excluding T1D (J) and in non-diabetic donors only (ND-Lean and ND-Obese) (K). **(L-M)** Fraction of immune cells relative to total nuclei versus age in all donors excluding T1D donors (L) and in non-diabetic donors only (ND-Lean and ND-Obese) (M). In (J-M) each dot represents a donor. Statistical significance was assessed using Spearman correlation analysis.

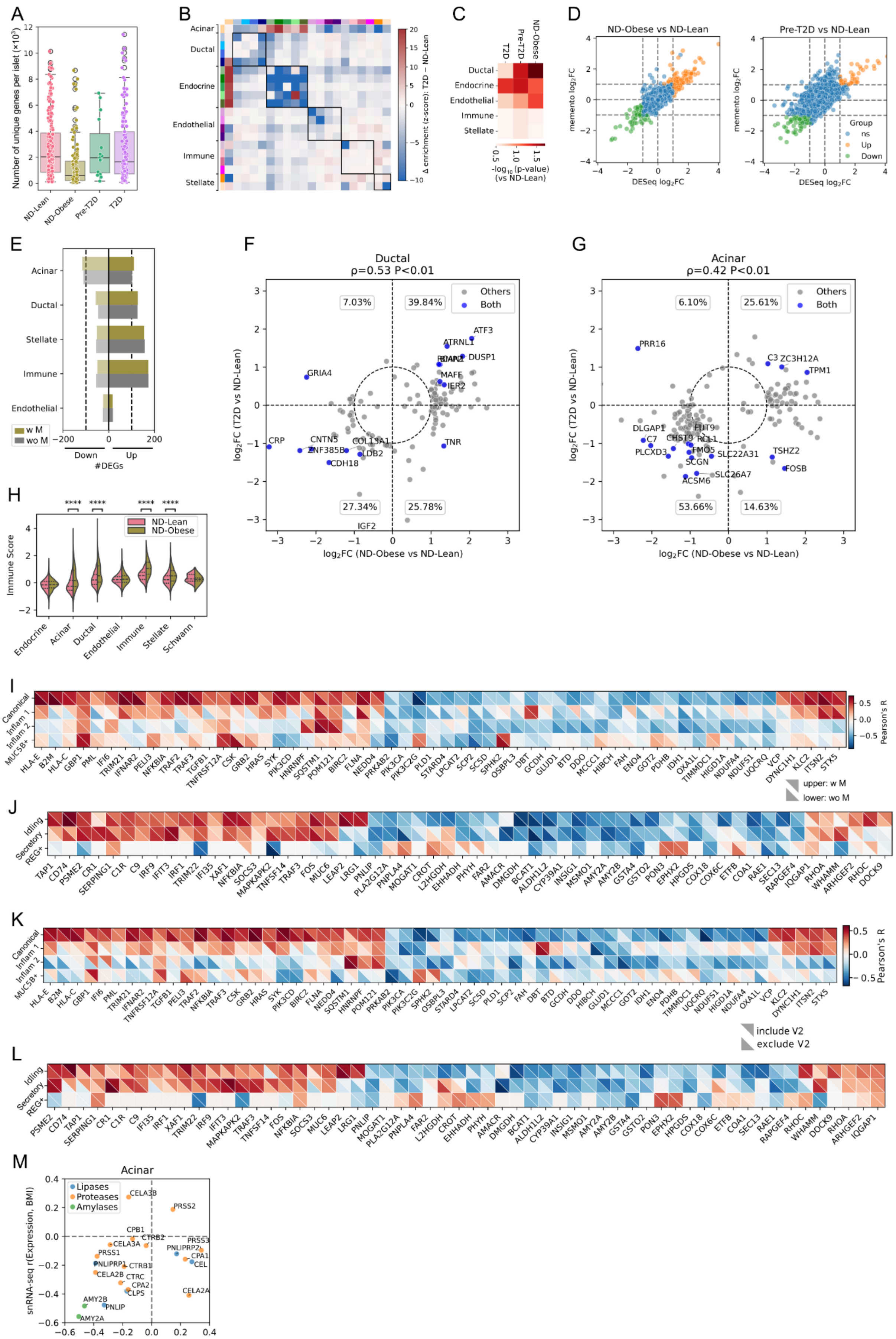
A
Number of unique genes per islet (×10³)
ND-Lean
ND-Obese
Pre-T2D
T2D
B
Acinar
Ductal
Endocrine
Endothelial
Immune
Stellate
Δ enrichment (z-score): T2D − ND-Lean
C
T2D
Pre-T2D
ND-Obese
Ductal
Endocrine
Endothelial
Immune
Stellate
-log10 (p-value) (vs ND-Lean)
D
ND-Obese vs ND-Lean
Pre-T2D vs ND-Lean
memento log2FC
DESeq log2FC
Group
ns
Up
Down
E
Acinar
Ductal
Stellate
Immune
Endothelial
w M
wo M
Down
Up
#DEGs
F
Ductal
ρ=0.53 P<0.01
Others
Both
7.03%
39.84%
27.34%
25.78%
log2FC (T2D vs ND-Lean)
log2FC (ND-Obese vs ND-Lean)
G
Acinar
ρ=0.42 P<0.01
6.10%
25.61%
53.66%
14.63%
H
Immune Score
ND-Lean
ND-Obese
Endocrine
Acinar
Ductal
Endothelial
Immune
Stellate
Schwann
I
Canonical
Inflam 1
Inflam 2
MUC5B+
Pearson's R
upper: w M
lower: wo M
J
Idling
Secretory
REG+
K
include V2
exclude V2
L
M
Acinar
Lipases
Proteases
Amylases
snRNA-seq r(Expression, BMI)
ST r(Expression, BMI)

**Supplementary Figure 2. Analysis of spatial transcriptomic data and exocrine correlations with BMI. (A)** Number of unique genes detected across donor groups. Each dot represents one islet. **(B)** Differential neighbourhood enrichment scores between T2D and ND-Lean samples. **(C)** Heatmap showing statistical significance (p-values) of changes in neighbourhood enrichment scores across pancreatic compartments in ND-Obese, Pre-T2D, and T2D samples relative to ND-Lean controls. P-values were calculated using permutation tests in which neighbourhood enrichment scores were randomly shuffled (Frobenius statistics, 10,000 permutations). **(D)** Comparison of log2 fold changes estimated by memento and DESeq in acinar cells (snRNA-seq) for ND-Obese versus ND-Lean (left) and Pre-T2D versus ND-Lean (right). (**E**) Number of differentially expressed genes (DEGs) in each cell type for ND-Obese status compared to ND-Lean based on snRNA-seq data. (**F-G**) Log2 fold changes between i) ND-Obese and ND-Lean (exclude male donors) or ii) T2D and ND-Lean (include male donors) for DEGs identified in **Fig. 2E-F** in ductal (F) and acinar cells (G). (**H**) Enrichment scores of immune system-associated genes identified by pathway analysis across all cell types in ND-Obese compared with ND-Lean (exclude male donors). Statistical significance was assessed using the Mann–Whitney test with Bonferroni-adjusted p-values (* $p < 0.05$, ** $p < 0.01$, and *** $p < 0.001$). (**I-J**) Heatmaps showing genes with high Pearson's correlation coefficients between pseudo-bulk expression (snRNA-seq) and BMI in ductal (I) and acinar (J) subtypes from non-diabetic female donors. For each gene, the upper right triangle includes all non-diabetic donors, whereas the lower left triangle excludes male donors. **(K-L)** Heatmaps showing genes with high Pearson's correlation coefficients between pseudo-bulk expression (snRNA-seq) and BMI in ductal (K) and acinar (L) subtypes from non-diabetic donors. For each gene, the upper right triangle includes all non-diabetic donors, whereas the lower left triangle excludes the donor with the highest BMI (V2, BMI=42). (**M**) Pearson correlation coefficients between gene expression and BMI for digestive enzyme genes in acinar cells from non-diabetic samples in spatial transcriptomics (ST) and snRNA-seq.

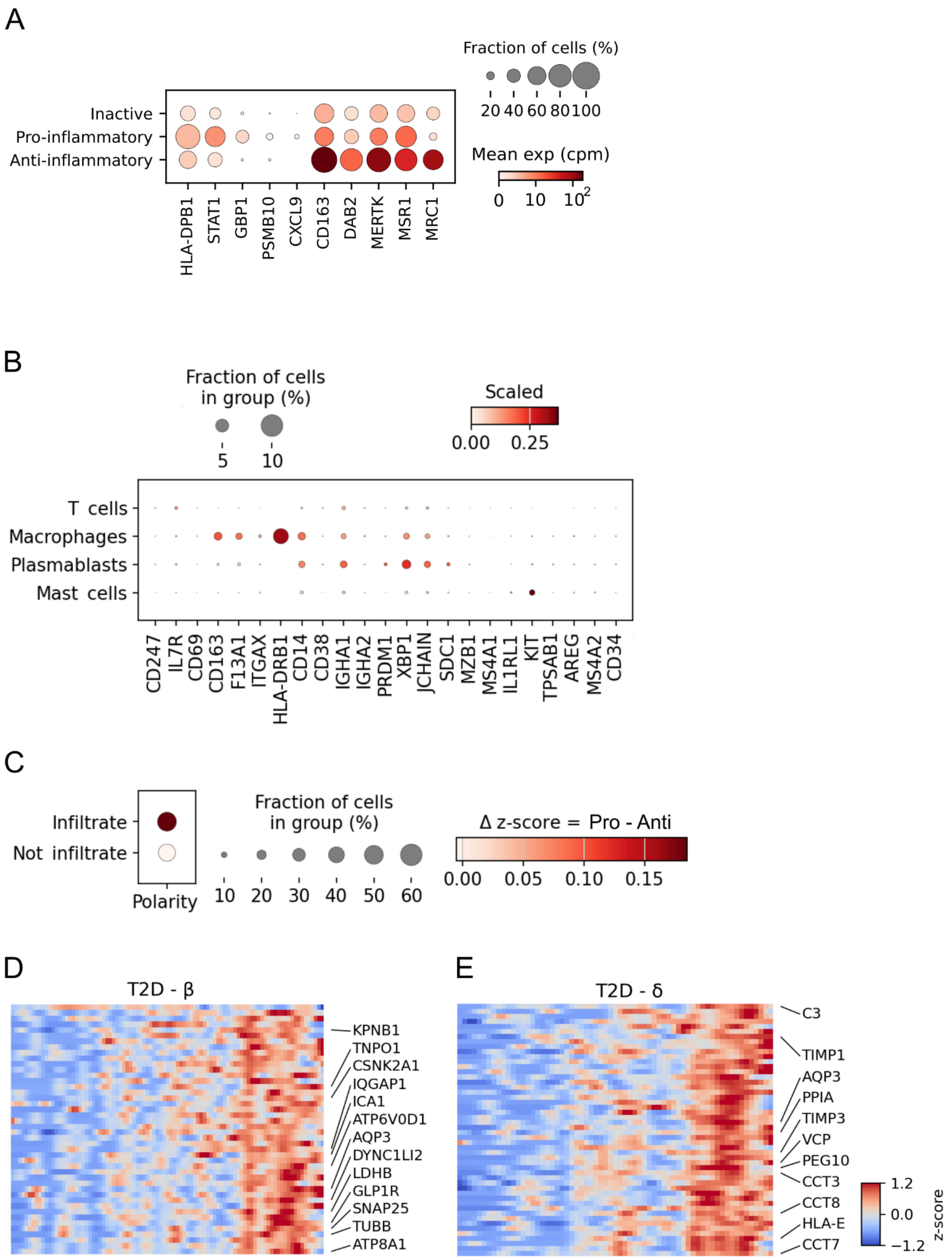


**Supplementary Figure 3. Analysis of macrophage polarization and macrophage-associated transcriptional changes in T2D islets.** **(A)** Dot plot showing normalized expression of marker genes in pro- and anti-inflammatory macrophages in snRNA-seq. **(B)** Dot plot showing scaled expression of selected immune marker genes across cell types in spatial transcriptomics. **(C)** Polarity score comparing pro-inflammatory and anti-inflammatory macrophage states in islet-infiltrating and non-infiltrating macrophages. Infiltrating macrophages are defined as those located ≤ 20 µm from an islet. **(D–E)** Heatmaps showing genes positively associated with macrophage infiltration in β cells (D) and δ cells (E) in islets from T2D donors.

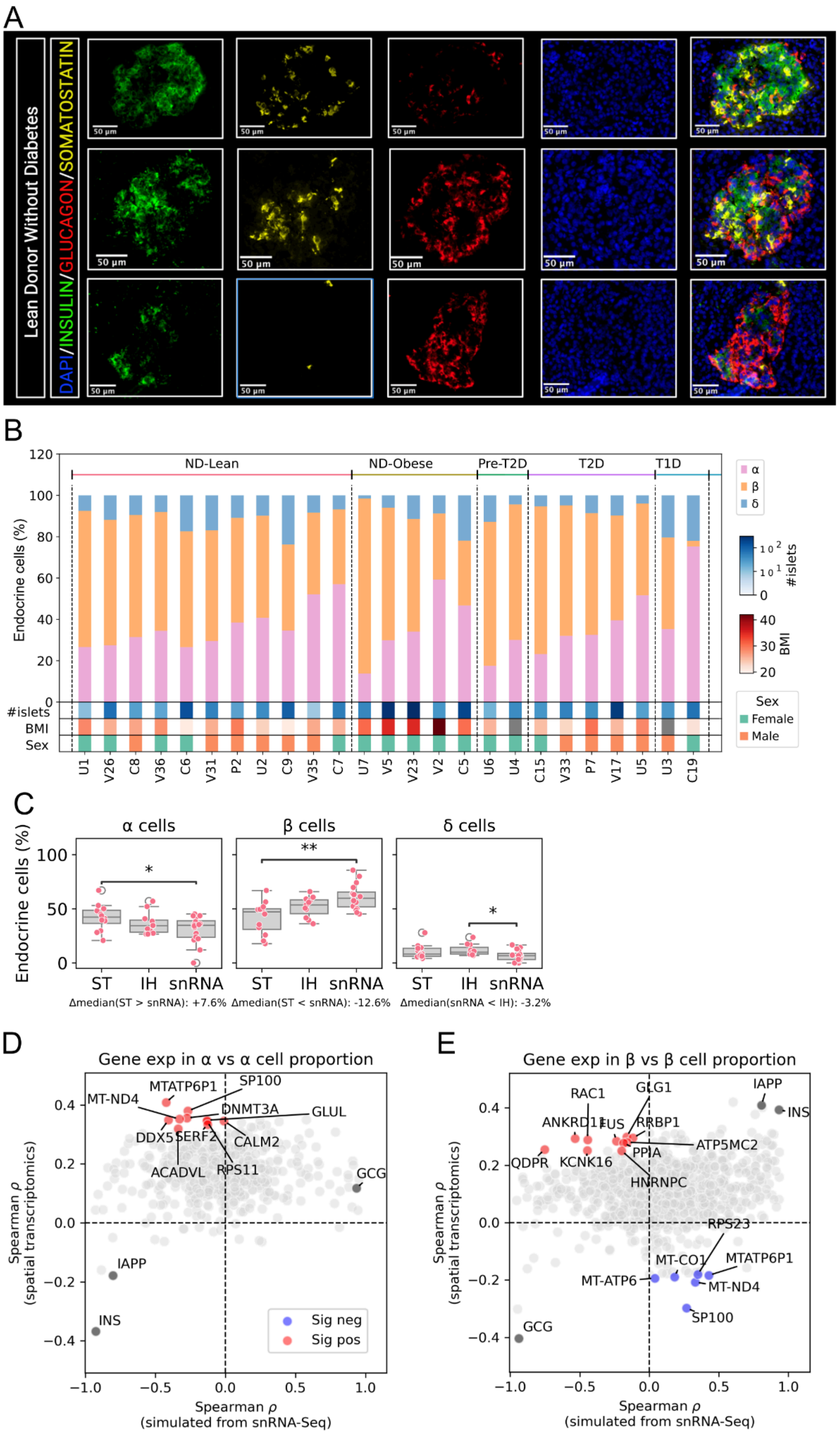
A
Lean Donor Without Diabetes
DAPI/INSULIN/GLUCAGON/SOMATOSTATIN
50 μm
B
ND-Lean
ND-Obese
Pre-T2D
T2D
T1D
Endocrine cells (%)
#islets
BMI
Sex
Female
Male
α
β
δ
U1 V26 C8 V36 C6 V31 P2 U2 C9 V35 C7 U7 V5 V23 V2 C5 U6 U4 C15 V33 P7 V17 U5 U3 C19
C
α cells
β cells
δ cells
Endocrine cells (%)
ST
IH
snRNA
Δmedian(ST > snRNA): +7.6%
Δmedian(ST < snRNA): -12.6%
Δmedian(snRNA < IH): -3.2%
D
Gene exp in α vs α cell proportion
MTATP6P1
SP100
MT-ND4
DNMT3A
GLUL
DDX5
SERF2
CALM2
ACADVL
RPS11
GCG
IAPP
INS
Sig neg
Sig pos
Spearman ρ (spatial transcriptomics)
Spearman ρ (simulated from snRNA-Seq)
E
Gene exp in β vs β cell proportion
RAC1
GLG1
IAPP
INS
ANKRD11
FUS
RRBP1
ATP5MC2
QDPR
KCNK16
PPIA
HNRNPC
RPS23
MT-CO1
MTATP6P1
MT-ATP6
MT-ND4
SP100
GCG
Spearman ρ (spatial transcriptomics)
Spearman ρ (simulated from snRNA-Seq)

**Supplementary Figure 4. Analysis of islet cell type composition across modalities and gene expression associations from spatial transcriptomics. (A)** Individual fluorescence channels from immunostaining images of islets shown **in Fig. 4A**. **(B)** Proportions of α, β, and δ cells measured by immunostaining for each donor. Lower annotations indicate the number of analyzed islets per donor, BMI, and sex. **(C)** Comparison of α, β, and δ cell proportions measured by spatial transcriptomics (ST), immunohistochemistry (IH), and snRNA-seq among ND-Lean samples. Each dot represents one donor. $^{*}p < 0.05$, $^{**}p < 0.01$ (Mann–Whitney U test). **(D–E)** Scatter plots showing Spearman's correlation coefficients between gene expression and α cell (D) or β cell (E) proportion in ND-Lean islets (spatial transcriptomics) against simulated islets generated from snRNA-seq data. Genes in the first and third quadrants show concordant associations between simulated and spatial data, likely driven by composition effects, whereas genes in the second and fourth quadrants show associations not captured by the simulations.

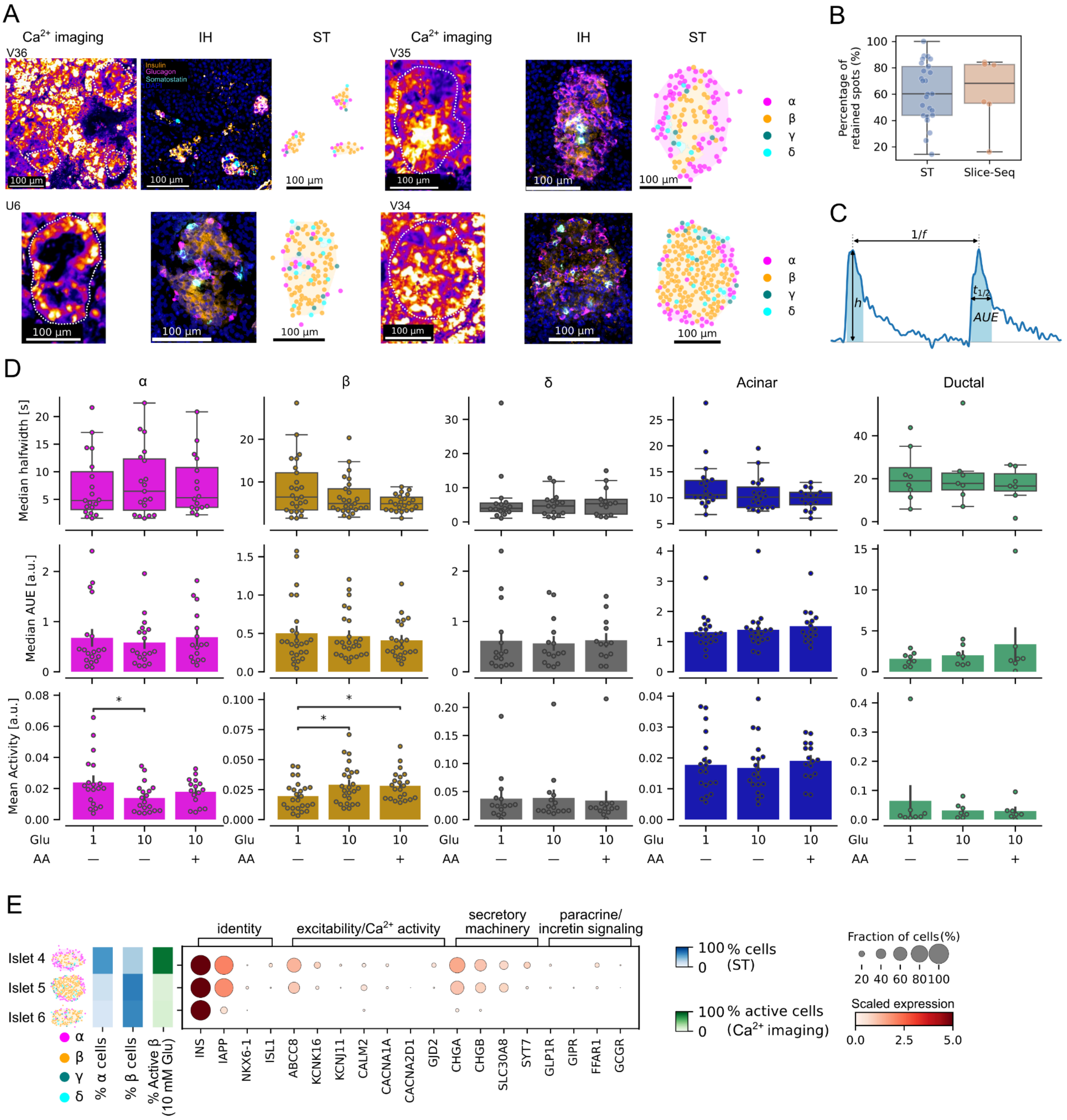


**Supplementary Figure 5. Analysis of Slice-seq spatial transcriptomics and calcium imaging data. (A)** Calcium imaging (left; dashed lines indicate islet boundaries), immunostaining (middle), and spatial transcriptomics (right) in Slice-seq islets (**Supp. Video 2-5**). **(B)** Percentage of retained spatial transcriptomics spots in samples analyzed with Slice-seq compared to standard spatial transcriptomics (ST). Each dot represents one tissue slice. **(C)** Schematic illustrating calcium event features, including event height (*h*), half-width ($t_{1/2}$), frequency (1/*f*) and area under the event (*AUE*), used for calcium trace analysis. **(D)** Median half-width, median AUE, and activities (i.e., area under the whole trace) across pancreatic cell types in acute tissue slices from ND-Lean donors under different glucose and amino acid (AA) conditions; α (N=237 in 20 islets), β (N=648 in 26 islets), δ (N=59 in 15 islets), acinar (N=2070) and ductal (N=94) cells in 7 ND-Lean donors. *$p < 0.05$, **$p < 0.01$ (Mann–Whitney U test with Bonferroni correction) **(E)** Gene expression in three large Slice-seq islets with different cell type composition and activity. From left to right: islet cell type composition, β cell activity (10 mM Glucose), and β cell pseudobulk expression (within the islet) of selected regulators of β cell excitability, secretory machinery, and paracrine signaling.

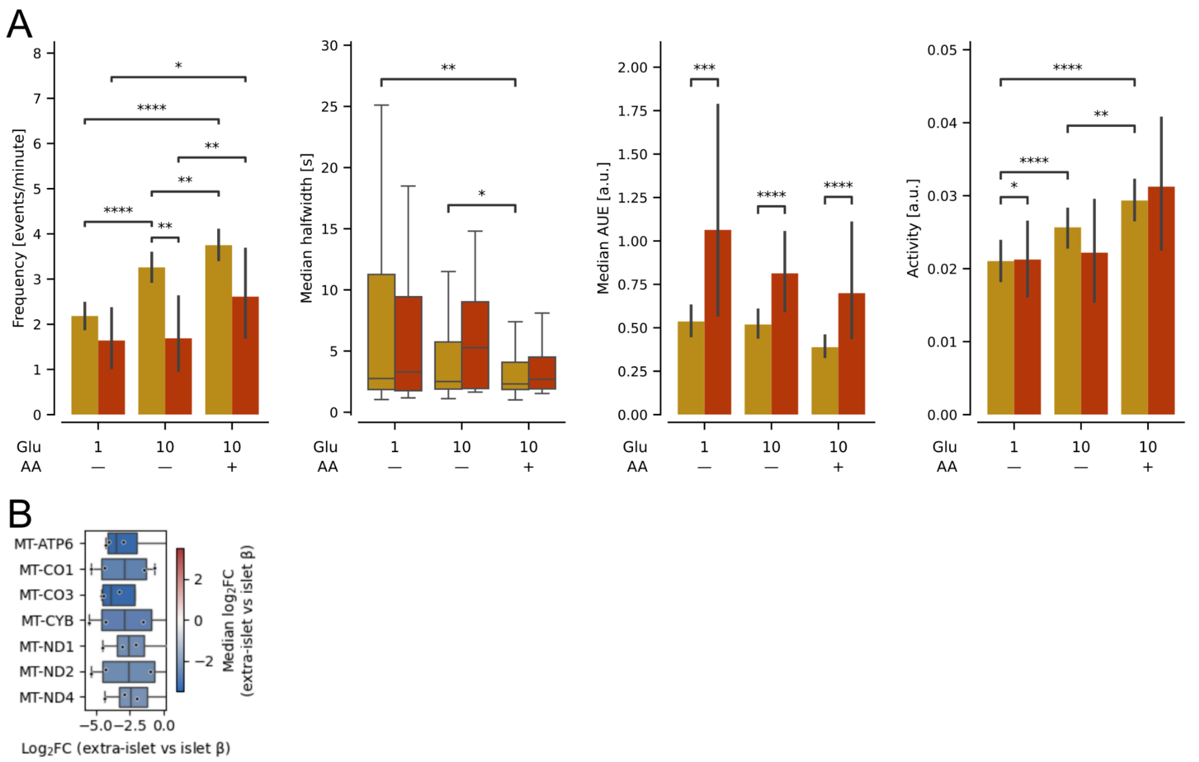


**Supplementary Figure 6. Calcium activity parameters in islet and extra-islet β cells.** (**A**) Frequency of calcium events, median half-width, median AUE, and activity in islet and extra-islet β cells under different glucose and amino acid (AA) conditions; extra-islet cells N=45, islet β cells N=648 in 7 ND-Lean donors. $^{*}p < 0.05$, $^{**}p < 0.01$, $^{***}p < 0.001$, $^{****}p < 0.0001$ (Mann–Whitney U test with Bonferroni correction). (**B**) Top differentially expressed genes between extra-islet and islet β cells based on spatial transcriptomics. Each dot represents a Slice-seq donor average.

A

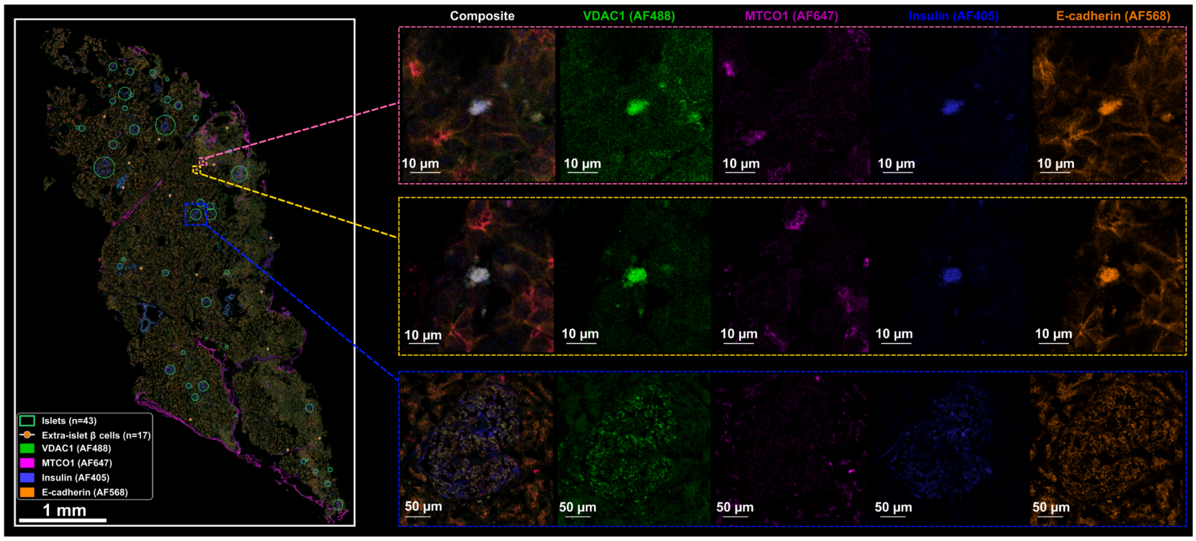

**Supplementary Figure 7. Segmentation of islet and extra-islet β cells.** (**A**) Immunofluorescence image showing segmentation of islets (green circles) and extra-islet β cells (yellow dots) based on insulin and E-cadherin staining. Insets show representative extra-islet β cells and an islet with individual fluorescence channels for VDAC1, MTCO1, insulin, and E-cadherin.
