## Supplementary Table for "Spatial and functional mapping of the human pancreas reveals endocrine and exocrine cell states in health and metabolic disease"

| Donor ID | BMI | Diabetes Duration | HbA1c (%) | Age | Sex | Part of the Pancreas | Doantion Type | Group | Treatment | Cohort ID | Cohort | Method |
|---|---|---|---|---|---|---|---|---|---|---|---|---|
| V55 | 18 | 0 | 5.50% | 62 | f | Tail | NDD | ND-lean | | | Vienna | Calcium imaging |
| C11 | 19.5 | 0 | 6.50% | 69 | f | Tail | NDD | ND-lean | | R552 | Edmonton | snRNA-Seq |
| V56 | 20 | 0 | 5.70% | 69 | f | Tail | NDD | ND-lean | | | Vienna | Calcium imaging |
| C9 | 20.1 | 0 | 4.10% | 31 | m | Tail | NDD | ND-lean | | R550 | Edmonton | snRNA-Seq,IH |
| P1 | 20.2 | 0 | n/a | 88 | f | Neck | DCD | ND-lean | | 26/13 | Pisa | snRNA-Seq,IH,ST |
| C6 | 20.2 | 0 | 5.40% | 54 | f | Tail | DCD | ND-lean | | R543 | Edmonton | snRNA-Seq,IH,ST |
| V34 | 23 | 0 | n/a | 46 | f | Tail | NDD | ND-lean | | | Vienna | IH,Calcium imaging,Slice-seq |
| U2 | 23.4 | 0 | 5.40% | 72 | m | Body | n/a | ND-lean | | H2673 | Uppsala | snRNA-Seq,IH,ST |
| V58 | 24 | 0 | 5.60% | 50 | m | Tail | NDD | ND-lean | | | Vienna | Calcium imaging |
| P4 | 24.2 | 0 | n/a | 61 | m | Neck | DCD | ND-lean | | 26/62 | Pisa | snRNA-Seq |
| C7 | 25.5 | 0 | 5.10% | 25 | f | Tail | NDD | ND-lean | | R548 | Edmonton | snRNA-Seq,IH,ST |
| C12 | 25.7 | 0 | 5.60% | 44 | m | Tail | DCD | ND-lean | | R553 | Edmonton | snRNA-Seq |
| V26 | 26 | 0 | n/a | 51 | f | Tail | NDD | ND-lean | | | Vienna | snRNA-Seq,IH,ST |
| V31 | 26 | 0 | n/a | 58 | m | Tail | NDD | ND-lean | | | Vienna | snRNA-Seq,IH,ST |
| C10 | 26 | 0 | 4.30% | 57 | f | Tail | NDD | ND-lean | | R551 | Edmonton | snRNA-Seq |
| V35 | 26.1 | 0 | 5.40% | 41 | m | Tail | NDD | ND-lean | | | Vienna | IH,Calcium imaging,Slice-seq |
| C8 | 26.5 | 0 | 5.10% | 54 | m | Tail | DCD | ND-lean | | R549 | Edmonton | snRNA-Seq,IH,ST |
| P3 | 27.7 | 0 | n/a | 59 | f | Neck | DCD | ND-lean | | 26/24 | Pisa | snRNA-Seq |
| V9 | 28 | 0 | n/a | 48 | m | Tail | NDD | ND-lean | | | Vienna | snRNA-Seq,IH,ST |
| V36 | 28 | 0 | n/a | 77 | f | Tail | NDD | ND-lean | | | Vienna | IH,Calcium imaging,Slice-seq |
| U1 | 28.5 | 0 | 5.60% | 62 | f | Body | n/a | ND-lean | | H2671 | Uppsala | snRNA-Seq,IH,ST |
| C14 | 28.9 | 0 | 5.40% | 35 | m | Tail | NDD | ND-lean | | R555 | Edmonton | snRNA-Seq |
| P2 | 29.4 | 0 | n/a | 68 | m | Neck | DCD | ND-lean | | 26/16 | Pisa | snRNA-Seq,IH,ST |
| C5 | 30.3 | 0 | 4.50% | 23 | f | Tail | NDD | ND-obese | | R554 | Edmonton | snRNA-Seq,IH,ST |
| U7 | 30.5 | 0 | 5.40% | 72 | f | Body | n/a | ND-obese | | H2584 | Uppsala | snRNA-Seq,IH,ST |
| V5 | 35 | 0 | n/a | 59 | f | Tail | DCD | ND-obese | | | Vienna | snRNA-Seq,IH,ST |
| V23 | 35 | 0 | n/a | 48 | f | Tail | NDD | ND-obese | | | Vienna | snRNA-Seq,IH,ST |
| V2 | 42 | 0 | n/a | 29 | f | Tail | NDD | ND-obese | | | Vienna | snRNA-Seq,IH,ST |
| U6 | 25.5 | 0 | 6% | 62 | f | Body | n/a | Pre-T2D | | H2692 | Uppsala | snRNA-Seq,IH,ST,Calcium imaging,Slice-seq |
| U9 | 31.1 | 0 | 5.80% | 69 | m | Body | n/a | Pre-T2D | | H2603 | Uppsala | snRNA-Seq,IH,ST |
| U4 | n/a | 0 | 5.90% | 61 | f | Body | n/a | Pre-T2D | | H2684 | Uppsala | snRNA-Seq,IH,ST |
| V33 | 23 | n/a | n/a | 65 | m | Tail | NDD | T2D | Metformin, Simvastatin | | Vienna | snRNA-Seq,IH,ST |
| C15 | 24.9 | n/a | 6.70% | 67 | f | Tail | NDD | T2D | diet controlled | R557 | Edmonton | snRNA-Seq,IH,ST |
| V17 | 25 | n/a | n/a | 68 | m | Tail | NDD | T2D | | | Vienna | snRNA-Seq,IH,ST |
| U5 | 28.6 | n/a | 8.20% | 54 | m | Body | n/a | T2D | | H2686 | Uppsala | snRNA-Seq,IH,ST |
| P5 | 29.3 | n/a | n/a | 78 | f | Neck | DCD | T2D | Insulin | 26/14 | Pisa | snRNA-Seq,IH,ST |
| P7 | 29.9 | n/a | n/a | 74 | m | Neck | DCD | T2D | Metformin/Insulin | 26/74 | Pisa | snRNA-Seq,IH,ST |
| P6 | 31.1 | n/a | n/a | 80 | f | Neck | NDD | T2D | Insulin | 26/68 | Pisa | snRNA-Seq,IH,ST |
| C13 | 33.2 | 15 years | 6.80% | 66 | m | Tail | NDD | T2D | Jardiance, Metoprolo | R556 | Edmonton | snRNA-Seq,IH,ST |
| C19 | 21.8 | n/a | 5.50% | 32 | f | Tail | NDD | T1D | | R572 | Edmonton | IH,ST |
| U3 | n/a | 10 | 6.60% | 42 | m | Body | n/a | T1D | | H2685 | Uppsala | snRNA-Seq,IH,ST,Calcium imaging |
| U11 | 30.8 | >20 years | 8.60% | 67 | f | Body | n/a | T1D | | H2727 | Uppsala | IH,Calcium imaging,Slice-seq |
